# Ccrk-Mak/Ick kinase signaling axis is a ciliary transport regulator essential for retinal photoreceptor maintenance

**DOI:** 10.1101/2024.05.24.595694

**Authors:** Taro Chaya, Yamato Maeda, Ryotaro Tsutsumi, Makoto Ando, Yujie Ma, Naoko Kajimura, Teruyuki Tanaka, Takahisa Furukawa

## Abstract

Primary cilia are microtubule-based sensory organelles whose dysfunction leads to ciliopathies in humans. The formation, function, and maintenance of primary cilia depend crucially on intraflagellar transport (IFT); however, the regulatory mechanisms of IFT and their physiological roles are poorly understood. In the current study, we screened and identified that the ciliopathy kinase Mak is a ciliary tip-localized IFT regulator that cooperatively acts with the ciliopathy kinase Ick, an IFT regulator. Simultaneous disruption of *Mak* and *Ick* resulted in loss of photoreceptor ciliary axonemes and severe degeneration in the mouse retina. Mak overexpression restored ciliary defects caused by *Ick* deficiency in cultured cells. Gene delivery of *Ick* and pharmacological inhibition of FGF receptors, negative regulators of Ick, ameliorated retinal degeneration in *Mak^−/−^*mice. In addition, we identified that Ccrk kinase is an upstream activator of Mak and Ick in retinal photoreceptor cells. Furthermore, overexpression of Mak, Ick, and Ccrk and pharmacological inhibition of FGF receptors suppressed ciliopathy-related phenotypes caused by cytoplasmic dynein inhibition in cultured cells. Collectively, our results show that the Ccrk-Mak/Ick axis is an essential IFT regulator crucial for retinal photoreceptor maintenance. This study sheds light on pathological mechanisms underlying retinitis pigmentosa caused by mutations in the human *MAK* gene and presents activation of Ick as a potential therapeutic approach for this retinal degenerative disease.

## Introduction

Primary cilia are hair-like microtubule-based structures that extend from the basal bodies of almost all cell types, and play key roles in a variety of sensory functions across species (1, 2). A diverse range of signaling receptors, ion channels, and their downstream effectors localized to the primary cilia sense and decode extracellular stimuli, including light and Hedgehog morphogens (3). For example, retinal photoreceptor cells possess a light-sensory structure containing components of the phototransduction cascade, including Rhodopsin and cone opsins, the outer segments, which are the specialized primary cilia that share common structural features with those of primary cilia in other cell types (4). Therefore, cilia are recognized as the signaling centers for multiple transduction pathways. In humans, ciliary dysfunction causes diseases known as ciliopathies, which are characterized by a broad spectrum of pathologies, including polydactyly, craniofacial abnormalities, brain malformation, intellectual disability, obesity, diabetes, polycystic kidney disease, anosmia, hearing loss, and retinal degeneration (5–7). Although many causative genes for ciliopathies have been identified, how genetic changes lead to the clinical phenotypes of ciliopathies with variable expressivity and severity is largely unknown (8).

The formation, length control, and function of cilia rely on intraflagellar transport (IFT), a bidirectional protein transport system coordinated by three large protein complexes, IFT-B, IFT-A, and BBSome, with molecular motors along the ciliary axonemal microtubules. They assemble into highly repetitive polymers called IFT trains, which move ciliary cargoes along the axoneme in both anterograde and retrograde directions (9–11). The three IFT complexes also function in the import and export of ciliary proteins (12, 13). IFT-B mediates anterograde transport from the base to the tip of the cilium driven by the kinesin-2 motor, whereas IFT-A mediates retrograde transport from the tip back to the base powered by the cytoplasmic dynein-2 motor (9, 12). The BBSome acts as a cargo adaptor that mediates the exit of membrane proteins from cilia (12, 14, 15). Recent studies have revealed detailed 3D structures of IFT-B, IFT-A, and BBSome (16–25). Mutations in genes encoding IFT components are known to cause ciliopathies, including Bardet-Biedl syndrome (BBS) (7). At the tip of the cilia, IFT trains disassemble and reassemble for turnaround and retrograde transport (26); however, the underlying regulatory mechanisms remain poorly understood (27).

We previously identified intestinal cell kinase (Ick), also known as ciliogenesis-associated kinase 1 (Cilk1), as a regulator of IFT turnaround at the ciliary tip (28, 29). *Chlamydomonas reinhardtii* (*Chlamydomonas*) LF4, *Tetrahymena* LF4A, *Leishmania mexicana* LmxMPK9, and *Caenorhabditis elegans* (*C. elegans*) DYF-5, orthologues of Ick, are also reported to be involved in the regulation of IFT as well as cilia/flagella length and formation (30–35). *Ick* deficiency leads to dysregulation of ciliary length, impaired Hedgehog signaling, and the accumulation of IFT-B, IFT-A, and BBSome components at the tips of cilia, whereas Ick overexpression induces the accumulation of IFT-B, but not IFT-A or BBSome components at ciliary tips (28, 36–39). *Ick*-deficient mice exhibit neonatal lethality, with developmental abnormalities observed in multiple organs and tissues, including the bone, lung, kidney, and inner ear (40). In humans, homozygous loss-of-function mutations in the *ICK* gene cause endocrine-cerebro-osteodysplasia (ECO) syndrome, an autosomal recessive ciliopathy showing neonatal lethality with various developmental defects involving the endocrine, cerebral, and skeletal systems, and short rib-polydactyly syndrome (SRPS), an autosomal recessive ciliopathy characterized by perinatal lethality with short ribs, shortened and hypoplastic long bones, polydactyly, and multiorgan system abnormalities (41–43). In addition, heterozygous variants of the human *ICK* gene are associated with juvenile myoclonic epilepsy (44). Collectively, Ick is currently recognized as a critical regulator of the IFT turnaround step at the ciliary tips (16, 27).

In the present study, we screened and found that male germ cell-associated kinase (Mak) regulates the IFT turnaround step at the ciliary tip. Mouse models revealed that *Mak* and *Ick* genetically interact in IFT regulation and play a crucial role in retinal photoreceptor maintenance, although *Ick* had a minor role. Mak overexpression restored ciliary defects in *Ick*-deficient cultured cells. Gene delivery of *Ick* and pharmacological inhibition of FGF receptors, negative regulators of Ick, ameliorated retinal degeneration in *Mak^−/−^* mice, a retinitis pigmentosa model. In addition, Cell cycle-related kinase (Ccrk) was identified as a critical upstream regulator of Mak/Ick in retinal photoreceptor cells. Furthermore, overexpression of Mak, Ick, and Ccrk and pharmacological inhibition of FGF receptors rescued the ciliopathy-related phenotypes resulting from cytoplasmic dynein inhibition. Taken together, this study identified a kinase signaling pathway regulating the IFT that plays an essential role in retinal photoreceptor maintenance.

## Results

### Ick plays a minor role in the regulation of IFT in retinal photoreceptor cells

To investigate the role of Ick in retinal photoreceptor development and maintenance, we mated *Ick* flox mice (28) with *Dkk3-Cre* mice (45), which predominantly expresses Cre recombinase in retinal progenitor cells, generated *Ick* conditional knockout (CKO) mice, and first analyzed *Ick* CKO mice at 1 month (1M). To observe the subcellular localization of Ick in retinal photoreceptor cells, immunohistochemical analysis was performed using an anti-Ick antibody. We found Ick signals in the distal regions of the photoreceptor ciliary axonemes in the control retina but not in the *Ick* CKO retina (Fig. S1A). To examine whether *Ick* deficiency affects the distribution of IFT components in photoreceptor cilia, we immunostained retinal sections using anti-IFT88 (an IFT-B component), anti-IFT140 (an IFT-A component), and anti-acetylated α-tubulin (Actub, a ciliary axoneme marker) antibodies, and observed no substantial differences between the control and *Ick* CKO retinas (Fig. S1B). We also performed immunohistochemical analyses using marker antibodies against Rhodopsin (rod outer segments), S-opsin (S-cone outer segments), and M-opsin (M-cone outer segments) and observed no substantial differences between the control and *Ick* CKO rod and cone outer segments (Fig. S1C). To evaluate the electrophysiological properties of the *Ick* CKO retina, electroretinograms (ERGs) of *Ick* CKO mice were measured under dark-adapted (scotopic) and light-adapted (photopic) conditions. Under scotopic conditions, the amplitude of a-waves and b-waves, originating mainly from the population activity of rod photoreceptor cells (a-waves) and rod bipolar cells (b-waves), was unaltered between the control and *Ick* CKO mice (Fig. S1D, E). Similar to scotopic ERG, the amplitudes of photopic a-waves and b-waves, which mainly reflect the population activity of cone photoreceptor cells (a-waves) and cone ON bipolar cells (b-waves), were comparable between the control and *Ick* CKO mice (Fig. S1D, E).

To investigate the effects of *Ick* deficiency on retinal photoreceptor cells at later stages, we performed histological analyses using retinal sections from *Ick* CKO mice at 6M. Toluidine blue staining showed that the outer nuclear layer (ONL) thickness decreased in the *Ick* CKO retina compared with that in the control retina from postnatal day 14 (P14), probably due to the reduced proliferation of retinal progenitor cells (Fig. S2A, B) (28). In contrast, the proportion of the ONL thickness to the inner retinal layer thickness decreased in *Ick* CKO mice compared with that in the control mice at 6M, but not at P14 or 1M, indicating that the ONL thickness decreased more in *Ick* CKO mice compared with that in the control mice at 6M (Fig. S2C). Immunohistochemical examination using marker antibodies against Rhodopsin, S-opsin, and M-opsin showed mislocalization of M-opsin in the inner part of the photoreceptor cells in the *Ick* CKO retina, although there were no obvious differences in the Rhodopsin and S-opsin signals between the control and *Ick* CKO retinas (Fig. S2D). We also performed an ERG analysis and found that scotopic a- and b-waves and photopic b-wave amplitudes in *Ick* CKO mice were lower than those in the control mice (Fig. S2E, F). These results suggest slowly progressive retinal degeneration in *Ick* CKO mice and a minor role of Ick in regulating IFT in retinal photoreceptor cells.

### Mak plays a major role in the regulation of IFT in retinal photoreceptor cells

Based on the above results, we hypothesized that serine-threonine kinase(s) other than Ick function as IFT regulators in the retinal photoreceptor cells. We focused on nine serine-threonine kinases close to Ick in the phylogenetic tree of the human kinome (46) and examined their subcellular localization using FLAG-tagged constructs (Fig. 1A). We observed that Mak and cyclin-dependent kinase-like 5 (Cdkl5), as well as Ick, localized to ciliary tips in cultured cells (Fig. 1B). To test whether overexpression of Mak and Cdkl5 affects IFT, we transfected plasmids encoding FLAG-tagged IFT57 (an IFT-B component) into NIH3T3 cells with the Mak-or Cdkl5-expressing construct (Fig. 1A). IFT57 was distributed along the ciliary axoneme without overexpression of Mak or Cdkl5. Similar to Ick, Mak, but not Cdkl5, overexpression induced IFT57 accumulation at the ciliary tips (Fig. 1C, D, and S3A), suggesting that Mak, rather than Cdkl5, is a candidate IFT regulator. To investigate the functional roles of Cdkl5 in retinal photoreceptor cells, we examined the tissue distribution of *Cdkl5* transcripts by RT-PCR analysis using mouse tissue cDNAs at 4 weeks (4wks) of age and observed that *Cdkl5* is ubiquitously expressed in various tissues, including the retina (Fig. S3B). To evaluate the effects of *Cdkl5* deficiency on retinal function, we performed an ERG analysis and found no obvious differences in the amplitudes of scotopic and photopic a- and b-waves between *Cdkl5^+/Y^* and *Cdkl5^−/Y^* mice at 1M and 3M (Fig. S3C), suggesting that *Cdkl5* is dispensable for retinal photoreceptor function and maintenance. In contrast, we previously reported that *Mak* is highly expressed in retinal photoreceptor cells, and that *Mak^−/−^*mice exhibit progressive retinal degeneration (47, 48). Subsequent analyses showed that mutations in the human *MAK* gene cause the retinal degenerative disease retinitis pigmentosa (49, 50). Together, our results and those of previous reports imply that Mak, rather than Cdkl5, functions as an IFT regulator in retinal photoreceptor cells.

**Figure 1.**
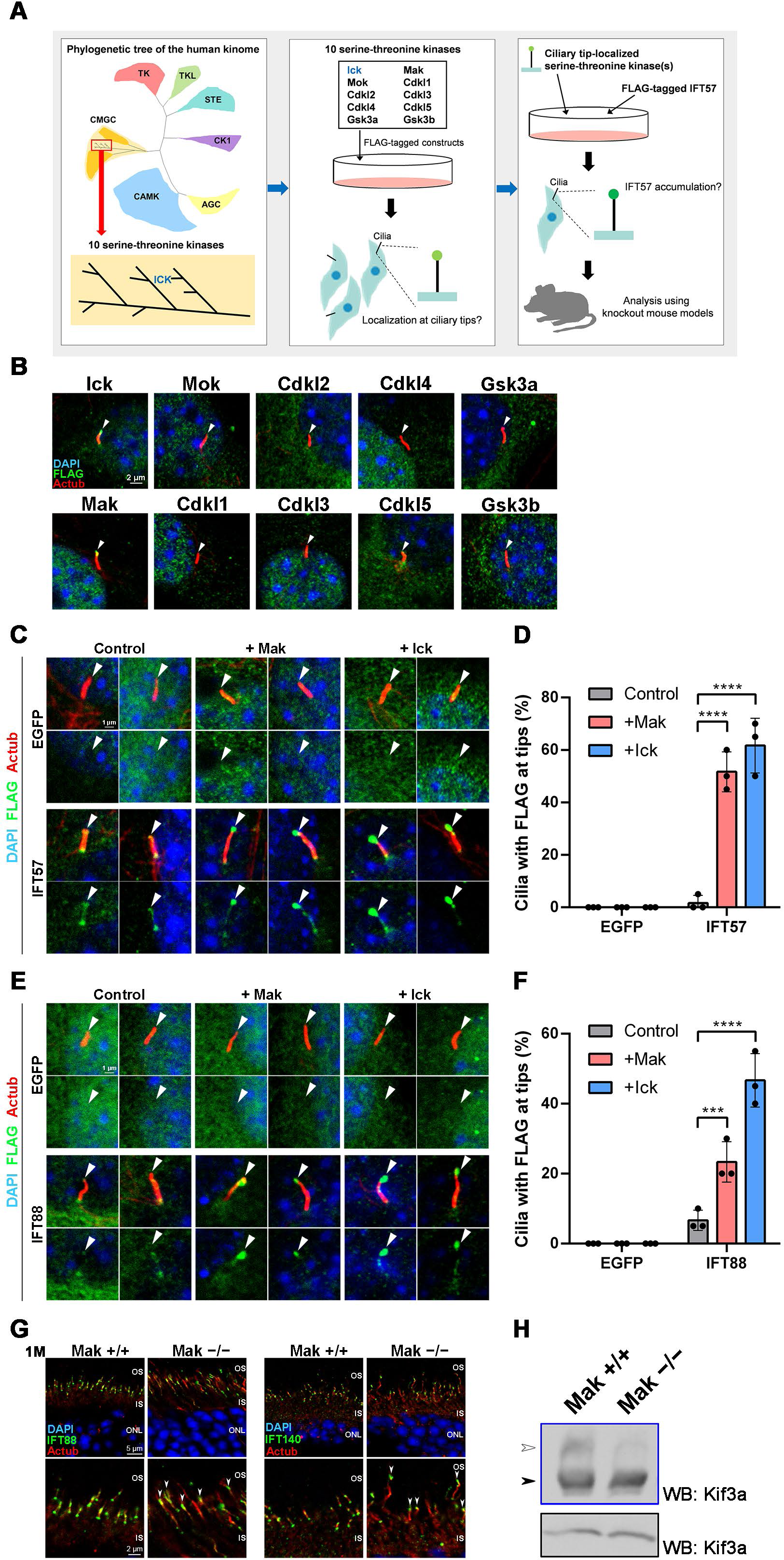
Ciliary localization of IFT components is changed by Mak overexpression or knockout. (A) Schematic diagrams of the experimental workflow for the screening of serine-threonine kinase(s) other than Ick that function as IFT regulators in retinal photoreceptor cells. (B) Subcellular localization of Ick and nine serine-threonine kinases that are close to Ick in the phylogenetic tree of the human kinome. A plasmid encoding a FLAG-tagged Ick, Mak, Mok, Cdkl1, Cdkl2, Cdkl3, Cdkl4, Cdkl5, Gsk3a, or Gsk3b was transfected into NIH3T3 cells. Cells were immunostained with anti-FLAG and anti-Actub antibodies. Arrowheads indicate ciliary tips. (C, D) Effects of Mak and Ick overexpression on ciliary localization of IFT57. (C) A FLAG-tagged EGFP or IFT57 expression plasmid was co-transfected into NIH3T3 cells with a plasmid expressing Mak or Ick. Cells were immunostained with anti-FLAG and anti-Actub antibodies. Arrowheads indicate ciliary tips. (D) The number of cilia with the FLAG-tagged EGFP or IFT57 predominantly localized at ciliary tips was counted. Data are presented as mean ± SD. \*\*\*\**p* < 0.0001 (two-way ANOVA followed by Tukey’s multiple comparisons test). n = 3 experiments. (E, F) Effects of Mak and Ick overexpression on ciliary localization of IFT88. (E) A FLAG-tagged EGFP or IFT88 expression plasmid was co-transfected into NIH3T3 cells with a plasmid expressing Mak or Ick. Cells were immunostained with anti-FLAG and anti-Actub antibodies. Arrowheads indicate ciliary tips. (F) The number of cilia with the FLAG-tagged EGFP or IFT88 predominantly localized at ciliary tips was counted. Data are presented as mean ± SD. \*\*\**p* < 0.001, \*\*\*\**p* < 0.0001 (two-way ANOVA followed by Tukey’s multiple comparisons test). n = 3 experiments. (G) Ciliary localization of IFT components in photoreceptor cells of the *Mak^−/−^* mouse retina. Retinal sections obtained from *Mak^+/+^* and *Mak^−/−^*mice at 1M were immunostained using antibodies against IFT88, IFT140, and Actub. IFT components were concentrated at the tips of photoreceptor ciliary axonemes in the *Mak^−/−^* retina (arrowheads). OS, outer segment; IS, inner segment; ONL, outer nuclear layer. (H) Kif3a phosphorylation in the *Mak^+/+^*and *Mak^−/−^* retina. The retinal lysates were analyzed by Phos-tag (blue box) and conventional (black box) Western blotting using an anti-Kif3a antibody. The level of Kif3a phosphorylation presented as the ratio of the upper band intensity (white arrowhead) to the lower band intensity (black arrowhead) decreased in the *Mak^−/−^*retina compared with that in the *Mak^+/+^* retina. Nuclei were stained with DAPI.

To assess the role of Mak in IFT in detail, we transfected plasmids encoding FLAG-tagged IFT88, IFT140, or BBS8 (a BBSome component) into NIH3T3 cells with or without the Mak-expressing construct. IFT88 was distributed along the ciliary axoneme in the absence of Mak overexpression. Under these conditions, IFT140 and BBS8 were localized mainly to the ciliary bases. Similar to Ick, Mak overexpression induced accumulation of IFT88, but not IFT140 or BBS8, at the ciliary tips (Fig. 1E, F, and S3D, E). The IFT140 signals at the ciliary bases increased in Mak- and Ick-overexpressing cells (Fig. S3D). We also observed the accumulation of IFT57 at the ciliary tips in human MAK- and ICK-overexpressing cells (Fig. S3F, G). To examine whether *Mak* deficiency affects the distribution of IFT components in retinal photoreceptor cilia, we immunostained retinal sections from *Mak^−/−^* mice at 1M using anti-IFT88, anti-IFT140, and anti-Actub antibodies. As observed in our previous study (47), photoreceptor ciliary axonemes were elongated in the *Mak^−/−^* retina compared to those in the *Mak^+/+^*retina (Fig. 1G). In contrast to the *Ick* CKO retinas (Fig. S1B), we observed that both IFT88 and IFT140 are concentrated at the tips of photoreceptor ciliary axonemes in the *Mak^−/−^* retina compared with those in the *Mak^+/+^*retina (Fig. 1G), which is similar to our previous observation that IFT components are concentrated at the ciliary tips in *Ick^−/−^* mouse embryonic fibroblasts (MEFs) (28). This observation shows that *Mak* deficiency as well as *Ick* deficiency impairs retrograde IFT. It has been previously reported that Ick phosphorylates Kif3a, a subunit of kinesin-2 (28, 51). To test whether Mak can also phosphorylate Kif3a, we performed Phos-tag Western blot analysis and observed a decrease in the up-shifted band of Kif3a in the *Mak^−/−^*retina compared to that in the *Mak^+/+^* retina, suggesting that Kif3a is phosphorylated by Mak in the retina (Fig. 1H). Together, these results suggest that Mak plays a major role in the regulation of IFT in retinal photoreceptor cells compared with Ick.

### Disruption of both *Mak* and *Ick* causes severe progressive retinal degeneration

To further investigate the functional roles of Mak and Ick in retinal photoreceptor cells, we generated *Mak Ick* double knockout (DKO) mice by mating *Mak^−/−^* mice with *Ick* CKO mice. We first performed histological analyses using retinal sections from *Mak Ick* DKO mice at P14 and 1M. Toluidine blue staining showed that the ONL thickness progressively decreased in *Mak^−/−^*and *Mak Ick* DKO retinas compared to that in the control retina, and that the extent of the decrease in the *Mak Ick* DKO retina was greater than that in the *Mak^−/−^* retina (Fig. 2A, B). Immunohistochemical examination using marker antibodies showed mislocalization of Rhodopsin, S-opsin, and M-opsin in the inner part of retinal photoreceptor cells in *Mak^−/−^*and *Mak Ick* DKO mice, as well as severe mislocalization in *Mak Ick* DKO mice compared to that in *Mak^−/−^* mice (Fig. 2C and S4A). We did not observe rod and cone outer segment structures in *Mak Ick* DKO mice (Fig. 2C and S4A). To examine whether both *Mak* and *Ick* deficiency affects cilia formation and the distribution of IFT components in the cilia of retinal photoreceptor cells, we immunostained retinal sections from *Mak Ick* DKO mice at P9 and P14 using antibodies against Actub, γ-tubulin (γTub, a basal body marker), pericentrin (a basal body marker), and IFT88. Notably, photoreceptor ciliary axonemes were not observed in the *Mak Ick* DKO retina, although those in the *Mak^−/−^* retina were elongated (Fig. 2D and S4B). We observed that IFT88 signals were concentrated near the basal bodies of retinal photoreceptor cells in *Mak Ick* DKO mice (Fig. 2D). To observe the ultrastructure of retinal photoreceptor cilia, we performed electron microscopic analysis. Although we found basal bodies, connecting cilia and outer segments were not observed in the retina of *Mak Ick* DKO mice at P14, which is consistent with the results of immunohistochemical analysis (Fig. 2E, F). To evaluate the electrophysiological properties of the *Mak Ick* DKO retina, we performed an ERG analysis and found that scotopic and photopic a- and b-wave amplitudes in *Mak^−/−^* mice were lower than those in the control mice (Fig. 2G, H). In *Mak Ick* DKO mice, no significant ERG responses were detected (Fig. 2G, H). These results suggest that deletion of both *Mak* and *Ick* causes severe retinal degeneration compared to deletion of *Mak* alone, and that *Mak* and *Ick* genetically interact and play a central role in the regulation of IFT in retinal photoreceptor cells.

**Figure 2.**
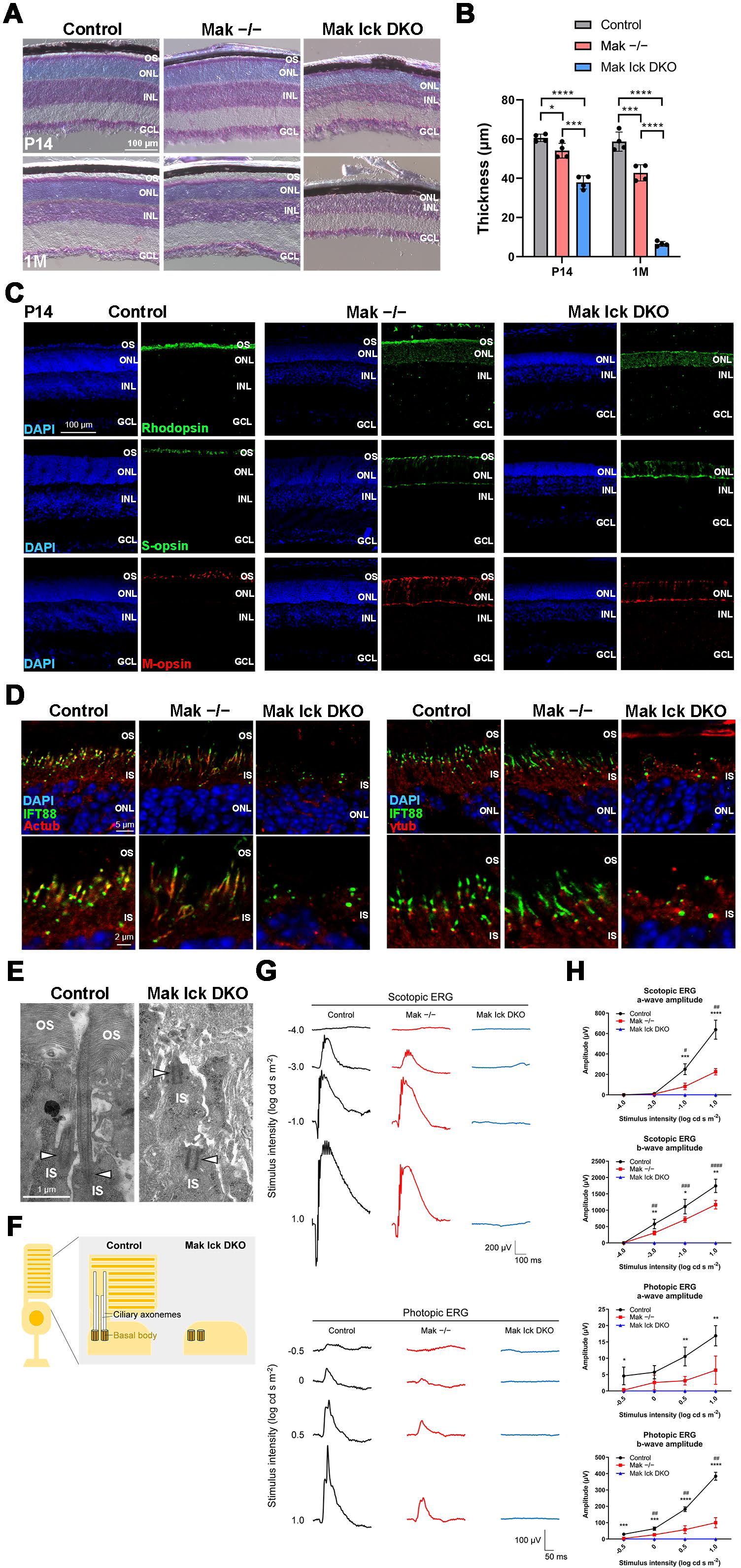
Severe progressive retinal degeneration in *Mak Ick* DKO mice. (A, B) Toluidine blue staining of retinal sections from the control, *Mak^−/−^*, and *Mak Ick* DKO mice at P14 and 1M. The ONL thickness was measured. Data are presented as mean ± SD. \**p* < 0.05, \*\*\**p* < 0.001, \*\*\*\**p* < 0.0001 (one-way ANOVA followed by Tukey’s multiple comparisons test). n = 4 mice per each genotype. (C) Immunostaining of retinal sections from the control, *Mak^−/−^*, and *Mak Ick* DKO mice at P14 using marker antibodies against Rhodopsin, S-opsin, and M-opsin. Severe photoreceptor outer segment disorganization and mislocalization of Rhodopsin and cone opsins were observed in the *Mak Ick* DKO retina. (D) Ciliary localization of IFT components in photoreceptor cells of the *Mak Ick* DKO retina. Retinal sections obtained from the control, *Mak^−/−^*, and *Mak Ick* DKO mice at P14 were immunostained using antibodies against IFT88, Actub, and γ-tubulin (γtub) (a marker for basal bodies). The ciliary axoneme of retinal photoreceptor cells were absent in *Mak Ick* DKO mice. (E) Longitudinal profiles of the connecting cilia in the P14 control and *Mak Ick* DKO photoreceptors observed by electron microscopy. Arrowheads indicate basal bodies. Connecting cilia were absent in the *Mak Ick* DKO retina. (F) Schematic representation of retinal photoreceptor cilia in the control and *Mak Ick* DKO mice. Although basal bodies were observed, ciliary axonemes and outer segments were not observed in the *Mak Ick* DKO retina. (G, H) ERG analysis of *Mak Ick* DKO mice. (G) Representative scotopic and photopic ERGs elicited by four different stimulus intensities (−4.0 to 1.0 log cd s/m^2^ and −0.5 to 1.0 log cd s/m^2^, respectively) from the control, *Mak^−/−^*, and *Mak Ick* DKO mice at 1M. (H) The scotopic and photopic amplitudes of a- and b-waves are shown as a function of the stimulus intensity. Data are presented as mean ± SD. One-way ANOVA followed by Tukey’s multiple comparisons test, Control vs. *Mak^−/−^* indicated by asterisks, \**p* < 0.05, \*\**p* < 0.01, \*\*\**p* < 0.001, \*\*\*\**p* < 0.0001, *Mak^−/−^*vs. *Mak Ick* DKO indicated by hash symbols, ^#^*p* < 0.05, ^##^*p* < 0.01, ^###^*p* < 0.001, ^####^*p* < 0.0001, n = 4, 4, and 3 mice (control, *Mak^−/−^*, and *Mak Ick* DKO, respectively). Nuclei were stained with DAPI. OS, outer segment; IS, inner segment; ONL, outer nuclear layer; INL, inner nuclear layer; GCL, ganglion cell layer.

### Heterozygous deletion of *Ick* exacerbates retinal degeneration in *Mak*^−/−^ mice

Previous reports have shown that loss-of-function mutations and variants are present in the human *ICK* gene (41–44). To gain insight into the effects of heterozygous *ICK* mutations or variants on the symptoms of retinitis pigmentosa caused by homozygous mutations in *MAK* gene in humans, we sought to generate and analyze *Mak^−/−^*; *Ick^+/−^*mice. We analyzed a publicly available single-cell RNA-sequencing (scRNA-seq) dataset of adult human retinas (52). In total, 20 cell clusters were identified using a principal component analysis (PCA)-based approach and projected by a uniform manifold approximation and projection (UMAP) onto a two-dimensional plot (Fig. 3A). We observed the expression of *MAK* and *ICK* in photoreceptor cells of the human retina (Fig. 3A, B). We confirmed that *MAK* and *ICK* are expressed in the human retina by RT-PCR analysis (Fig. S5A). To investigate the effects of *Ick* heterozygous deficiency on retinal photoreceptor cells, we performed histological analyses using retinal sections from *Mak^+/−^*; *Ick^+/−^* mice at 6M. We previously observed no obvious retinal structural differences between wild-type and *Mak^+/−^* mice at 6M (47). Toluidine blue staining showed no significant differences in the ONL thickness between the *Mak^+/+^*; *Ick^+/+^* and *Mak^+/−^*; *Ick^+/−^* retinas (Fig. S5B, C). Immunohistochemical examination using marker antibodies against Rhodopsin, S-opsin, and M-opsin showed no obvious differences between the *Mak^+/+^*; *Ick^+/+^* and *Mak^+/−^*; *Ick^+/−^* retinas (Fig. S5D). To evaluate the effects of *Ick* heterozygous deficiency on retinal function, we performed ERG analysis and found no significant differences in the amplitudes of scotopic and photopic a- and b-waves between *Mak^+/+^*; *Ick^+/+^* and *Mak^+/−^*; *Ick^+/−^* mice (Fig. S5E, F). These results show that *Ick* heterozygous deficiency alone does not affect retinal photoreceptor function or maintenance.

**Figure 3.**
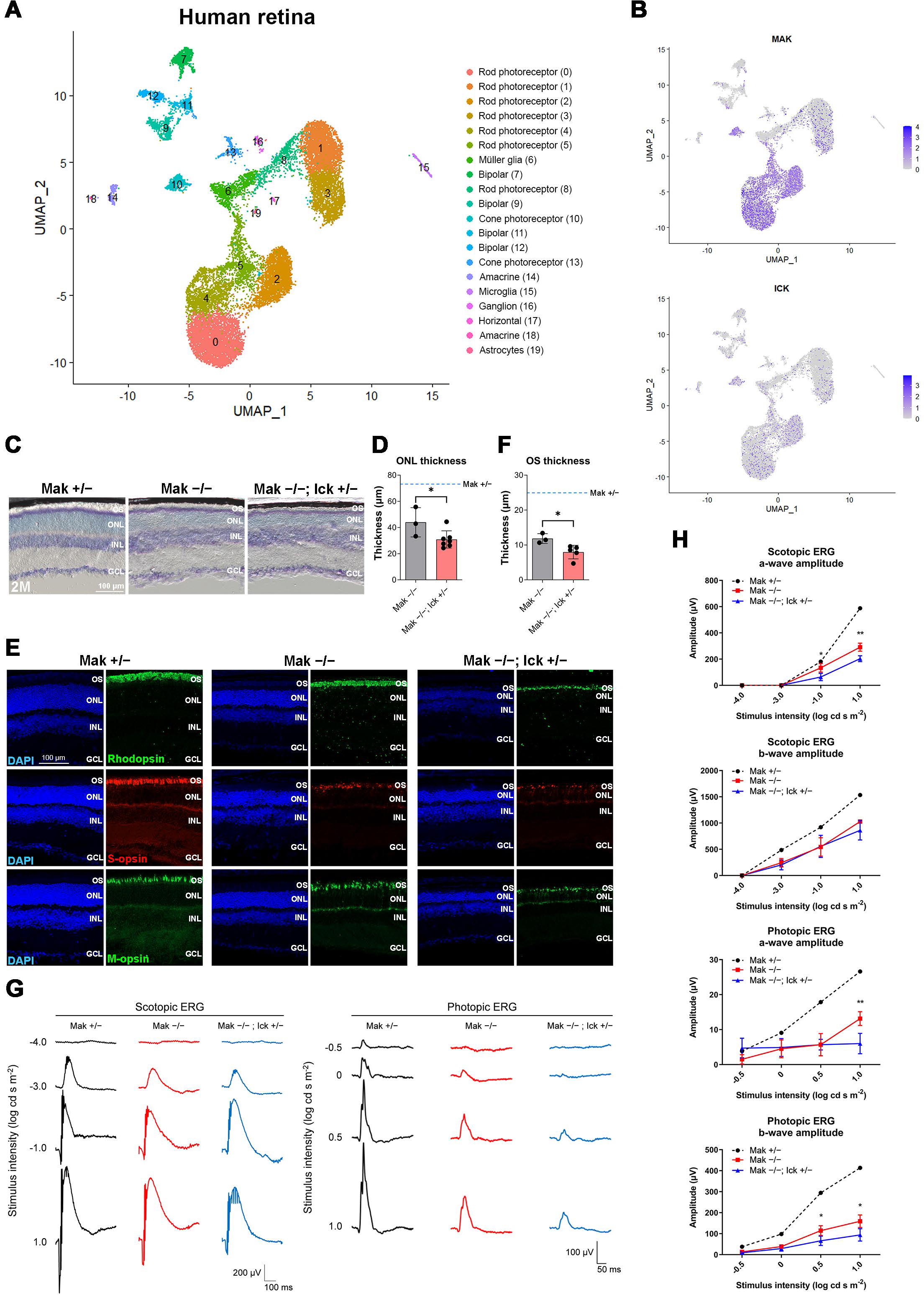
*Ick* haploinsufficiency exacerbates retinal degeneration caused by loss of *Mak*. (A) Uniform manifold approximation and projection (UMAP) visualization of retinal cells in adult human retinas. (B) Feature plots showing expression of *MAK* and *ICK* in retinal cells of adult human retinas. (C, D) Toluidine blue staining of retinal sections from *Mak^+/−^*, *Mak^−/−^*, and *Mak^−/−^*; *Ick^+/−^* mice at 2M. The ONL thickness was measured. Data are presented as mean ± SD. \**p* < 0.05 (unpaired t-test). n = 5, 3, and 7 mice (*Mak^+/−^*, *Mak^−/−^*, and *Mak^−/−^*; *Ick^+/−^*, respectively). The blue dotted line indicates the ONL thickness in the *Mak^+/−^* retina. (E, F) Immunostaining of retinal sections from *Mak^+/−^*, *Mak^−/−^*, and *Mak^−/−^*; *Ick^+/−^* mice at 2M using marker antibodies against Rhodopsin, S-opsin, and M-opsin. Nuclei were stained with DAPI. The rod outer segment length was measured. Data are presented as mean ± SD. \**p* < 0.05 (unpaired t-test). n = 4, 3, and 5 mice (*Mak^+/−^*, *Mak^−/−^*, and *Mak^−/−^*; *Ick^+/−^*, respectively). The blue dotted line indicates the rod outer segment length in the *Mak^+/−^* retina. (G, H) ERG analysis of *Mak^−/−^*; *Ick^+/−^* mice at 2M. (G) Representative scotopic and photopic ERGs elicited by four different stimulus intensities (−4.0 to 1.0 log cd s/m^2^ and −0.5 to 1.0 log cd s/m^2^, respectively) from *Mak^+/−^*, *Mak^−/−^*, and *Mak^−/−^*; *Ick^+/−^* mice. (H) The scotopic and photopic amplitudes of a- and b-waves are shown as a function of the stimulus intensity. Data are presented as mean ± SD. \**p* < 0.05, \*\**p* < 0.01 (unpaired t-test). n = 5, 3, and 6 mice (*Mak^+/−^*, *Mak^−/−^*, and *Mak^−/−^*; *Ick^+/−^*, respectively). OS, outer segment; ONL, outer nuclear layer; INL, inner nuclear layer; GCL, ganglion cell layer.

Next, we performed histological analyses of retinal sections from *Mak^−/−^*; *Ick^+/−^* mice at 2M. Toluidine blue staining showed that the ONL thickness decreased in *Mak^−/−^* and *Mak^−/−^*; *Ick^+/−^*retinas compared with that in *Mak^+/−^* retina, and that the extent of the decrease in *Mak^−/−^*; *Ick^+/−^* retina was greater than that in *Mak^−/−^* retina (Fig. 3C, D). Immunohistochemical examination using marker antibodies showed disorganized structures of the rod and cone outer segments in *Mak^−/−^* and *Mak^−/−^*; *Ick^+/−^* mice, and severe disorganization in *Mak^−/−^*; *Ick^+/−^* mice compared with that in *Mak^−/−^* mice (Fig. 3E). The rod outer segment length decreased in the *Mak^−/−^*; *Ick^+/−^* retina compared to that in the *Mak^−/−^* retina (Fig. 3E, F). To evaluate the electrophysiological properties of the *Mak^−/−^*; *Ick^+/−^* retina, we performed an ERG analysis and found that scotopic and photopic a- and b-wave amplitudes in *Mak^−/−^* and *Mak^−/−^*; *Ick^+/−^* mice were lower than those in *Mak^+/−^* mice, and that the amplitudes of scotopic a-waves and photopic a- and b-waves significantly decreased in *Mak^−/−^*; *Ick^+/−^* mice compared to those in *Mak^−/−^* mice (Fig. 3G, H). These results suggest that *ICK* can be a modifier of retinitis pigmentosa caused by *MAK* mutations in humans.

### The abnormalities caused by *Ick* and *Mak* deficiency are rescued by activation of Mak and Ick, respectively

To examine whether ciliary defects caused by *Ick* deficiency can be restored by Mak activation, we transfected plasmids encoding Mak into *Ick^−/−^* MEFs. We previously reported that ciliated cells are fewer and ciliary length is shorter in *Ick^−/−^* MEFs than in *Ick^+/+^* MEFs (28). Remarkably, the overexpression of Mak, similar to that of Ick, rescued the ciliary defects observed in *Ick^−/−^*MEFs (Fig. 4A-C). To investigate whether retinal abnormalities caused by *Mak* deficiency can also be restored by Ick activation, we first injected adeno-associated virus (AAV) expressing Ick under the control of the Rhodopsin kinase promoter (53) into the *Mak^−/−^* retina (Fig. 4D). We observed that the mislocalization of Rhodopsin to the inner part of rod photoreceptor cells in the *Mak^−/−^* retina was partially suppressed by Ick expression in retinal photoreceptor cells (Fig. 4E, F). It was previously reported that fibroblast growth factor receptors (Fgfrs) negatively regulate Ick activity (54). Next, we injected BGJ398, an FDA-approved small-molecule inhibitor of Fgfrs, into *Mak^−/−^* mice to activate Ick in retinal photoreceptor cells (Fig. 4G). Toluidine blue staining showed an increase in the ONL thickness of the retina in BGJ398-treated *Mak^−/−^* mice compared with that in vehicle-treated *Mak^−/−^* mice (Fig. 4H, I). To evaluate the effects of BGJ398 on retinal function in *Mak^−/−^* mice, we performed ERG analysis and observed that scotopic a- and b-wave amplitudes in BGJ398-treated *Mak^−/−^* mice were higher than those in vehicle-treated *Mak^−/−^* mice (Fig. 4J). To test the effects of BGJ398 on the activity of Fgfrs in the retina, we performed Phos-tag Western blot analysis and observed that the Ick substrate Kif3a was more phosphorylated in the retina of BGJ398-treated *Mak^−/−^*mice than in that of vehicle-treated *Mak^−/−^* mice, suggesting that Ick is activated by BGJ398 in the *Mak^−/−^* retina (Fig. 4K, L). To examine whether BGJ398-mediated rescue of the defects in the *Mak^−/−^* retina is through Ick, we injected BGJ398 into *Mak Ick* DKO mice (Fig. S6A). In contrast to the *Mak^−/−^*retina, toluidine blue staining showed no significant differences in the ONL thickness of the retinas between vehicle- and BGJ398-treated *Mak Ick* DKO mice (Fig. S6B, C). To evaluate the effects of BGJ398 on retinal function in *Mak Ick* DKO mice, we performed ERG analysis and observed no obvious differences in scotopic a- and b-wave amplitudes between vehicle- and BGJ398-treated *Mak Ick* DKO mice (Fig. S6D). These data from BGJ398-treated *Mak Ick* DKO mice support the idea that the rescue effects of BGJ398 on retinal degeneration in *Mak^−/−^* mice are mediated by Ick. To test the effects of BGJ398 on wild-type retinas, we injected BGJ398 into *Mak^+/+^* mice and performed histological and electrophysiological analyses (Fig. S6A). In contrast to *Mak^−/−^* mice, we observed no significant differences in the ONL thickness or scotopic ERG a- and b-wave amplitudes between vehicle- and BGJ398-treated *Mak^+/+^*mice (Fig. S6E-G). Collectively, these results show that Mak activation can rescue the abnormalities caused by *Ick* deficiency and *vice versa*.

**Figure 4.**
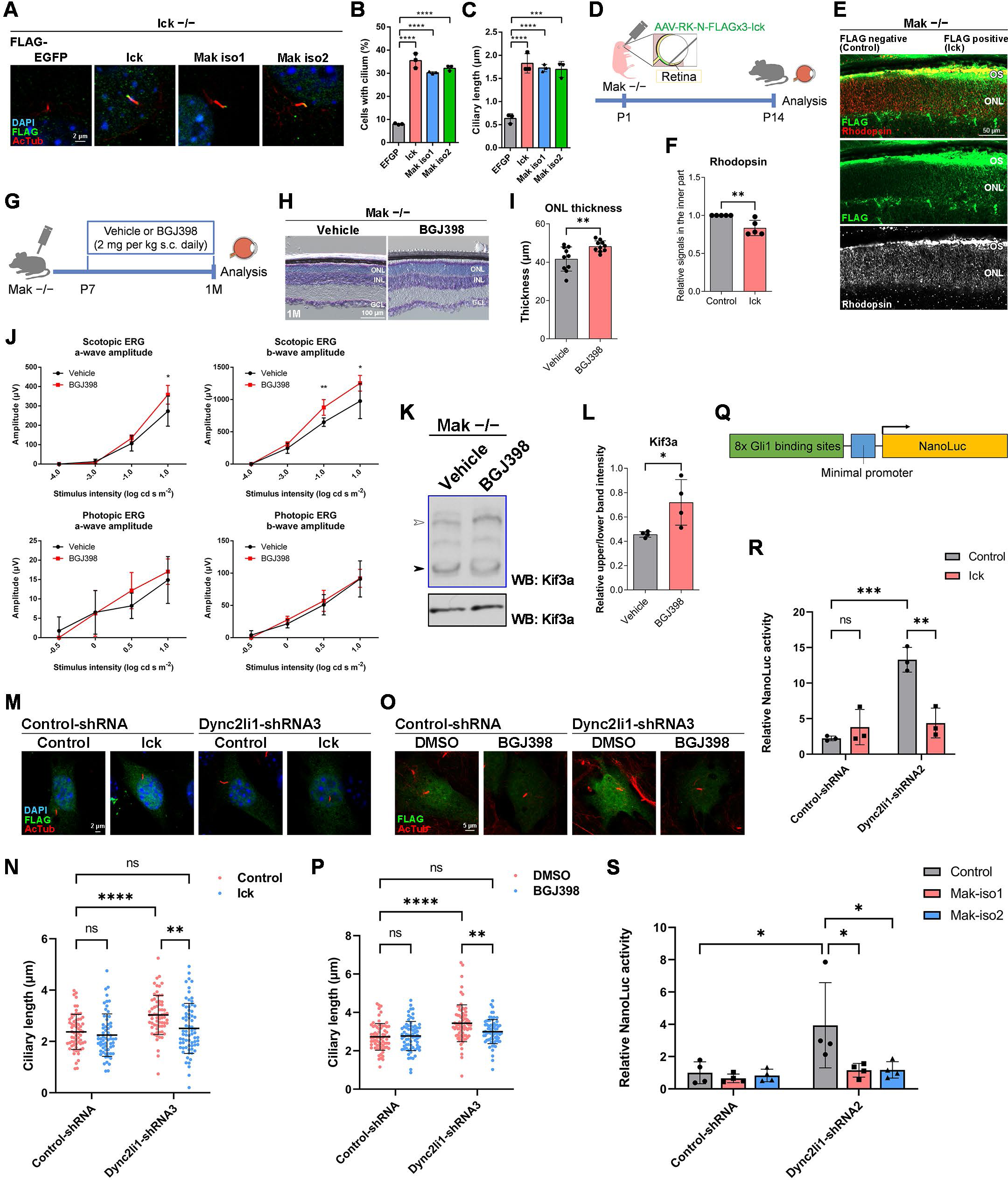
Activation of Mak and Ick can rescue the abnormalities caused by *Ick*, *Mak*, and *Dync2li1* deficiency. (A-C) Effects of Mak overexpression on ciliary defects in *Ick^−/−^* MEFs. (A) A plasmid encoding a FLAG-tagged EGFP, Ick, Mak isoform1 (Mak iso1), or Mak isoform2 (Mak iso2) was transfected into *Ick^−/−^* MEFs. Cells were immunostained with anti-FLAG and anti-Actub antibodies. The numbers (B) and length (C) of the cilia stained with an antibody against Actub in FLAG-positive cells were measured. Data are presented as mean ± SD. \*\*\**p* < 0.001, \*\*\*\**p* < 0.0001 (one-way ANOVA followed by Tukey’s multiple comparisons test). n = 3 experiments. (D-F) Subcellular localization of Rhodopsin in Ick-overexpressing photoreceptor cells of the *Mak^−/−^* mouse retina. (D) Schematic diagram of schedule for AAV subretinal injection and harvest of retinas. AAV expressing FLAG-tagged Ick driven by the Rhodopsin kinase promoter was injected into P1 *Mak^−/−^* mouse retinas. At P14, their retinas were harvested, sectioned, and immunostained with anti-FLAG and anti-Rhodopsin antibodies. (E) The Rhodopsin signals in the inner part of photoreceptors decreased in FLAG-positive regions. (F) The immunofluorescence signals of Rhodopsin detected in the inner part of photoreceptors were quantified using ImageJ software. The signals of Rhodopsin in the inner part of photoreceptors were normalized to the total (OS + the inner part) signals of Rhodopsin in photoreceptors. Rhodopsin signals in the inner part (normalized to the total signals) of FLAG-positive regions relative to those of FLAG-negative regions were then calculated. Data are presented as mean ± SD. \*\**p* < 0.01 (unpaired t-test). n = 5 retinas from four mice. (G) Schematic diagram of schedule for drug administration. BGJ398, an inhibitor of Fgfrs, was injected into *Mak^−/−^* mice from P7 to 1M every day. s.c., subcutaneous. (H, I) Toluidine blue staining of retinal sections from 1M *Mak^−/−^* mice treated with or without BGJ398, which was injected into the mice from P7 to 1M every day. Data are presented as mean ± SD. \*\**p* < 0.01 (unpaired t-test). n = 10 mice each. (J) ERG analysis of 1M *Mak^−/−^* mice treated with or without BGJ398, which was injected into the mice from P7 to 1M every day. The scotopic and photopic amplitudes of a- and b-waves are shown as a function of the stimulus intensity (−4.0 to 1.0 log cd s/m^2^ and −0.5 to 1.0 log cd s/m^2^, respectively). Data are presented as mean ± SD. \**p* < 0.05, \*\**p* < 0.01 (unpaired t-test). n = 6 mice each (scotopic), n = 5 and 6 mice (photopic) (vehicle-treated and BGJ398-treated, respectively). (K, L) Kif3a phosphorylation in the retina of 1M *Mak^−/−^* mice treated with or without BGJ398, which was injected into the mice from P7 to 1M every day. (K) The retinal lysates were analyzed by Phos-tag (blue box) and conventional (black box) Western blotting with the anti-Kif3a antibody. (L) Relative phosphorylation of Kif3a was quantified by the ratio of the upper band intensity (white arrowhead) to the lower band intensity (black arrowhead). Data are presented as mean ± SD. \**p* < 0.05 (unpaired t-test). n = 4 mice each. (M, N) Effects of Ick overexpression on ciliary length in cells knocked down for *Dync2li1*. (M) A plasmid encoding Control-shRNA or Dync2li1-shRNA3 was co-transfected into NIH3T3 cells with or without a plasmid expressing Ick in combination with a construct encoding FLAG-tagged EGFP. Cells were immunostained with anti-FLAG and anti-Actub antibodies. (N) The length of cilia stained with an antibody against Actub in FLAG-positive cells was measured. Data are presented as mean ± SD. \*\**p* < 0.01, \*\*\*\**p* < 0.0001, ns, not significant (two-way ANOVA followed by Tukey’s multiple comparisons test). Control-shRNA; Control, Control-shRNA; Ick, Dync2li1-shRNA3; Control, and Dync2li1-shRNA3; Ick, n = 66, 63, 67, and 70 cilia, respectively, from 3 experiments. (O, P) Effects of BGJ398 treatment on ciliary length in cells knocked down for *Dync2li1*. (O) NIH3T3 cells transfected with plasmids expressing FLAG-tagged EGFP and Control-shRNA or Dync2li1-shRNA3 were treated with 100 nM BGJ398 or DMSO for 24 h before harvest. Cells were immunostained with anti-FLAG and anti-Actub antibodies. (N) The length of cilia stained with an antibody against Actub in FLAG-positive cells was measured. Data are presented as mean ± SD. \*\**p* < 0.01, \*\*\*\**p* < 0.0001, ns, not significant (two-way ANOVA followed by Tukey’s multiple comparisons test). Control-shRNA; DMSO, Control-shRNA; BGJ398, Dync2li1-shRNA3; DMSO, and Dync2li1-shRNA3; BGJ398, n= 71, 70, 74, and 73 cilia, respectively, from 3 experiments. (Q-S) Luciferase reporter gene assay using 8x Gli1-binding sites-minimal promoter-NanoLuc luciferase constructs. (Q) Schematic diagram of the construct expressing NonoLuc luciferase under the control of 8x Gli1-binding sites and minimal promoter. NIH3T3 cells were transfected with plasmids expressing Dync2li1-shRNA2 and Ick (R), Mak iso1, or Mak iso2 (S) along with a Nanoluc luciferase reporter construct driven by the 8x Gli1-binding sites and minimal promoter and a Firefly luciferase-expressing construct driven by the SV40 promoter and enhancer. Luciferase activities of cell lysates were measured 24 h after serum starvation with 100 nM Smoothened agonist (SAG). Nanoluc luciferase activity was normalized to Firefly luciferase activity. Data are presented as mean ± SD. \**p* < 0.05, \*\**p* < 0.01, \*\*\**p* < 0.001, ns, not significant (two-way ANOVA followed by Tukey’s multiple comparisons test). n = 3 (R) and n = 4 (S) experiments. Nuclei were stained with DAPI. OS, outer segment; ONL, outer nuclear layer; INL, inner nuclear layer; GCL, ganglion cell layer.

### Ciliary defects resulting from cytoplasmic dynein inhibition are rescued by Mak and Ick activation

Given that deficiency of *Mak* and *Ick* impaired retrograde IFT, we hypothesized that Mak and Ick activation could also restore the ciliary abnormalities caused by retrograde IFT defects related to other gene mutations. To test this hypothesis, NIH3T3 cells were transfected with plasmids expressing Ick and treated with Ciliobrevin D, a cytoplasmic dynein inhibitor, and immunofluorescence analysis was performed to observe cilia. We found that the reduced number of cilia in cells treated with Ciliobrevin D is partially restored by Ick expression (Fig. S6H, I). Next, we performed knockdown and rescue experiments to observe cilia. We constructed shRNAs to knock down *Dync2li1*, a ciliopathy gene encoding cytoplasmic dynein-2 light intermediate chain 1, and confirmed that Dync2li1 expression levels decreased in cells expressing Dync2li1-shRNA2, Dync2li1-shRNA3, Dync2li1-shRNA5, and Dync2li1-shRNA6 (Fig. S6J). We transfected the Control-shRNA, Dync2li1-shRNA3, or Dync2li1-shRNA5 expression plasmids into NIH3T3 cells with plasmids encoding Ick and examined the cilia by immunostaining using an anti-Actub antibody. We found that the expression of Ick rescued Dync2li1-shRNA-induced cilia elongation (Fig. 4M, N, and S6K, L). We also transfected the Control-shRNA or Dync2li1-shRNA3 expression plasmids into NIH3T3 cells, treated the cells with BGJ398, and observed that Dync2li1-shRNA-mediated cilia elongation was suppressed by BGJ398 treatment (Fig. 4O, P). To examine whether aberrant Hedgehog signaling due to dynein inhibition is rescued by Ick and Mak activation, we performed a reporter gene assay using a NanoLuc luciferase reporter construct driven by 8x Gli1-binding sites and a minimal promoter (Fig. 4Q). Luciferase activity increased in cells expressing Dync2li1-shRNA2, Dync2li1-shRNA3, and Dync2li1-shRNA6 compared to that in cells expressing Control-shRNA, indicating that Hedgehog signaling was activated by *Dync2li1* knockdown (Fig. 4R, S, and S6M). We found that the expression of Ick and Mak suppressed *Dync2li1* knockdown-induced Hedgehog signaling activation (Fig. 4R, S). These results demonstrate that Ick and Mak activation can rescue ciliary defects caused by cytoplasmic dynein inhibition.

### *Ccrk* depletion causes severe progressive retinal degeneration very similar to that observed in *Mak Ick* DKO mice

To elucidate regulatory mechanisms of Mak and Ick activity in retinal photoreceptor cells, we focused on phosphorylation of the Mak and Ick proteins, since high-throughput phosphoproteomic analyses have shown that Mak and Ick are extensively phosphorylated at residue 157 (PhosphoSitePlus) (55), which is located in the kinase domain showing a high similarity of amino acid sequence and predicted structure between Mak and Ick (Fig. 5A). To examine whether this residue is phosphorylated in the retina, we performed a western blot analysis using an antibody against phosphorylated Mak Thr-157 (pMak) and detected pMak in the *Mak^+/+^* retina (Fig. 5B). To investigate the physiological role of Thr-157 phosphorylation, we generated a construct encoding the Ick protein harboring a Thr-to-Ala mutation at residue 157 (Ick-T157A). NIH3T3 cells were transfected with plasmids expressing EGFP, wild-type Ick (Ick-WT), Ick-T157A, or Ick harboring a Lys-to-Arg mutation at residue 33 (Ick-kinase dead (KD)), and immunofluorescence analysis was performed using an antibody against phosphorylated Kif3a Thr-674 (pKif3a). As previously observed (28), cytoplasmic pKif3a signals increased in cells expressing Ick-WT compared with those in cells expressing EGFP (Fig. 5C). In contrast, the expression of Ick-T157A and Ick-KD did not increase cytoplasmic pKif3a signals, suggesting that Thr-157 phosphorylation is required for Mak and Ick activity (Fig. 5C). To identify kinase(s) phosphorylating Thr-157, we searched for the serine-threonine kinases belonging to “Kinase, *in vitro*” and/or top 5 kinases listed in the “Kinase Prediction” tool (56) for phosphorylation of Mak and Ick Thr-157 using PhosphoSitePlus, and picked up Ccrk, Map3k5, Map3k15, Map2k2, Stk16, Mos, and Stk32b (Fig. 5D). Among the genes encoding them, a previous ChIP-seq analysis showed that the binding site of Crx, a master transcription factor in retinal photoreceptor maturation and survival (57, 58), is only assigned to the *Ccrk* gene, suggesting that *Ccrk* can be a candidate gene expressed in retinal photoreceptor cells (Fig. 5D) (59). We examined the tissue distribution of the transcripts of *Ccrk* and *Bromi*/*Tbc1d32*, which encodes a physical and functional interaction partner of Ccrk (60), by RT-PCR analysis using mouse tissue cDNAs and found that *Ccrk* and *Bromi* are ubiquitously expressed in various tissues, including the retina (Fig. S7A). UMAP plots of neuronal and glial cells from adult human retinas (52) showed the expression of *CCRK* and *BROMI* in the retinal photoreceptor cells (Fig. 3A and 5E). We confirmed that *CCRK* and *BROMI* are expressed in the human retina by RT-PCR analysis (Fig. S7B). To examine the expression pattern of *Ccrk* in the retina, we performed *in situ* hybridization analysis using mouse retinal sections and observed that *Ccrk* is expressed in the ONL (Fig. S7C). Based on these, we focused on Ccrk in subsequent experiments.

**Figure 5.**
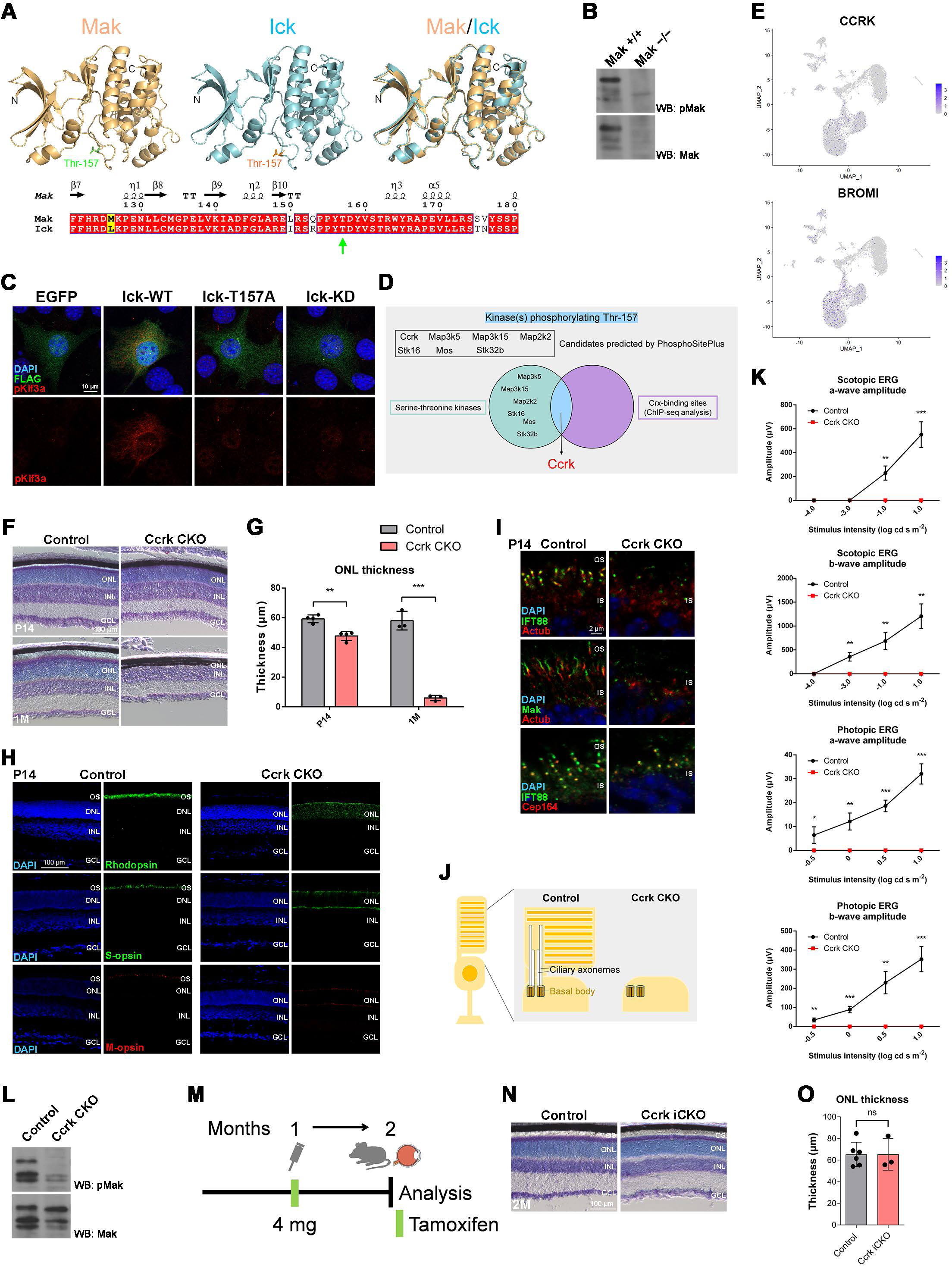
Severe progressive retinal degeneration in *Ccrk* CKO mice. (A) Thr-157 in the Mak and Ick proteins. (Upper panel) Predicted 3D structures of the kinase domains of the Mak and Ick proteins and a superposition of their structures are shown. (Lower panel) Structure-based sequence alignment of the Mak and Ick proteins prepared with ESPript. Similar and identical residues are marked by yellow and red boxes, respectively. Secondary structure assignment is based on the predicted structure of Mak. A green arrow indicates Thr-157. (B) Western blot analysis of the phosphorylated Mak Thr-157 (pMak) and Mak protein in *Mak^+/+^* and *Mak^−/−^* retinas. (C) Kif3a phosphorylation in Ick-overexpressing cells. A plasmid encoding a FLAG-tagged EGFP, wild-type Ick (Ick-WT), Ick harboring a Thr-to-Ala mutation at residue 157 (Ick-T157A), or Ick harboring a Lys-to-Arg mutation at residue 33 (Ick-kinase dead (KD)) was transfected into NIH3T3 cells. Cells were immunostained with anti-FLAG and anti-phosphorylated Kif3a Thr-674 (pKif3a) antibodies. Substantial cytoplasmic pKif3a signals were observed in cells expressing Ick-WT but not in those expressing EGFP, Ick-T157A, or Ick-KD. (D) Schematic diagram summarizing search for kinase(s) phosphorylating Mak and Ick Thr-157. (E) Feature plots showing expression of *CCRK* and *BROMI* in retinal cells of adult human retinas. (F, G) Toluidine blue staining of retinal sections from the control and *Ccrk* CKO mice at P14 and 1M. The ONL thickness was measured. Data are presented as mean ± SD. \*\**p* < 0.01, \*\*\**p* < 0.001 (unpaired t-test). n = 4 mice per each genotype at P14 and n = 3 mice per each genotype at 1M. (H) Immunostaining of retinal sections from the control and *Ccrk* CKO mice at P14 using marker antibodies against Rhodopsin, S-opsin, and M-opsin. Severe photoreceptor outer segment disorganization and mislocalization of Rhodopsin and cone opsins were observed in the *Ccrk* CKO retina. (I) Ciliary localization of IFT components in photoreceptor cells of the *Ccrk* CKO retina. Retinal sections obtained from the control and *Ccrk* CKO mice at P14 were immunostained using antibodies against IFT88, Actub, Mak, and Cep164 (a marker for basal bodies). The ciliary axoneme of retinal photoreceptor cells were absent in *Ccrk* CKO mice. (J) Schematic representation of retinal photoreceptor cilia in the control and *Ccrk* CKO mice. Although basal bodies were observed, ciliary axonemes and outer segments were not observed in the *Ccrk* CKO retina. (K) ERG analysis of *Ccrk* CKO mice. Scotopic and photopic ERGs elicited by four different stimulus intensities (−4.0 to 1.0 log cd s/m^2^ and −0.5 to 1.0 log cd s/m^2^, respectively) from the control and *Ccrk* CKO mice at 1M. The scotopic and photopic amplitudes of a- and b-waves are shown as a function of the stimulus intensity. Data are presented as mean ± SD. \**p* < 0.05, \*\**p* < 0.01, \*\*\**p* < 0.001 (unpaired t-test). n = 3 mice per each genotype. (L) Western blot analysis of pMak and the Mak protein in the control and *Ccrk* CKO retinas. (M) Schematic diagram of schedule for tamoxifen administration and analysis of mice. Mice were injected with tamoxifen at 1M and analyzed at 2M. (N, O) Toluidine blue staining of retinal sections from the control and *Ccrk* iCKO mice at 2M. The ONL thickness was measured. Data are presented as mean ± SD. ns, not significant (unpaired t-test). n = 6 and 3 mice (control and *Ccrk* iCKO, respectively). Nuclei were stained with DAPI. OS, outer segment; IS, inner segment; ONL, outer nuclear layer; INL, inner nuclear layer; GCL, ganglion cell layer.

To investigate the role of Ccrk in retinal photoreceptor development and maintenance, we generated *Ccrk* flox mice using targeted gene disruption (Fig. S7D). We mated *Ccrk* flox mice with *Dkk3-Cre* mice to generate *Ccrk* CKO mice (Fig. S7D). In the *Ccrk* CKO retina, no *Ccrk* mRNA or Ccrk protein expression was detected by RT-PCR and Western blotting with the anti-Ccrk antibody that we generated (Fig. S7E-G). We first performed histological analyses using retinal sections from *Ccrk* CKO mice at P14 and 1M. Toluidine blue staining showed a progressive decrease in the ONL thickness in the *Ccrk* CKO retinas compared to that in the control retinas (Fig. 5F, G). Immunohistochemical examination using marker antibodies showed mislocalization of Rhodopsin, S-opsin, and M-opsin in the inner part of the retinal photoreceptor cells in *Ccrk* CKO mice (Fig. 5H and S7H). We observed no obvious rod or cone outer segment structures in *Ccrk* CKO mice (Fig. 5H and S7H). To examine whether *Ccrk* deficiency affects cilia formation and the distribution of IFT components in the cilia of retinal photoreceptor cells, we immunostained retinal sections from *Ccrk* CKO mice at P14 using antibodies against Actub, Cep164 (a basal body marker), IFT88, and Mak. Similar to Ick, Mak was localized in the distal regions of the photoreceptor ciliary axonemes in the control retina (Fig. 5I). In contrast, photoreceptor ciliary axonemes were not observed in the *Ccrk* CKO retinas (Fig. 5I, J). We observed that IFT88 signals were concentrated near the basal bodies of retinal photoreceptor cells in *Ccrk* CKO mice (Fig. 5I). To evaluate the electrophysiological properties of the *Ccrk* CKO retina, we performed ERG analysis and observed no significant ERG responses in *Ccrk* CKO mice (Fig. 5K). The observed phenotypes of severe retinal degeneration in *Ccrk* CKO mice resemble those observed in *Mak Ick* DKO mice, suggesting that the Ccrk-Mak/Ick axis plays a crucial role in IFT regulation in retinal photoreceptor cells. To confirm this, we performed western blot analysis and found that pMak was markedly decreased in the *Ccrk* CKO retina compared with that in the control retina (Fig. 5L).

### Ccrk is dispensable for Mak phosphorylation maintenance *in vivo*

To investigate whether Ccrk is required for pMak maintenance *in vivo*, we mated *Ccrk* flox mice with *Crx-CreERT2* mice (61, 62), which express *CreERT2* predominantly in retinal photoreceptor cells, injected tamoxifen into *Ccrk^flox/flox^*; *Crx-CreERT2* mice at 1M, and analyzed the mice at 2M (Fig. 5M). RT-PCR and western blotting showed that *Ccrk* mRNA and Ccrk protein expression decreased in the retina of *Ccrk^flox/flox^*; *Crx-CreERT2* mice treated with tamoxifen (*Ccrk* iCKO mice) (Fig. S7I, J). Toluidine blue staining showed no significant differences in the ONL thickness between the control and *Ccrk* iCKO retinas (Fig. 5N, O). Immunohistochemical examination using marker antibodies against Rhodopsin, S-opsin, and M-opsin also showed no obvious differences between the control and *Ccrk* iCKO retinas (Fig. S7K). In addition, we performed an ERG analysis and found no significant differences in the amplitudes of scotopic and photopic ERG a- and b-waves between the control and *Ccrk* iCKO mice (Fig. S7L, M). Furthermore, we found no substantial differences in the amount of pMak between the control and *Ccrk* iCKO retinas (Fig. S7N). These results suggest that pMak is maintained by autophosphorylation (63) but not by Ccrk *in vivo*.

### *Ccrk* dosage affects retinal degeneration in *Mak^−/−^* mice and ciliary abnormalities caused by cytoplasmic dynein inhibition

To assess the relationship between Ccrk and Mak/Ick in retinal photoreceptor cells, we sought to generate and analyze *Mak^−/−^*; *Ccrk^+/−^* mice. To examine the effects of *Ccrk* heterozygous disruption on retinal photoreceptor cells, we compared the retinal phenotypes of *Mak^+/−^*; *Ccrk^+/−^* mice to those of *Mak^+/−^* mice at 2M. Immunohistochemical examination using marker antibodies against Rhodopsin, S-opsin, and M-opsin showed no obvious differences between the *Mak^+/−^* and *Mak^+/−^*; *Ccrk^+/−^*retinas (Fig. S8A). To evaluate the effects of *Ccrk* heterozygous deficiency on retinal function, we performed an ERG analysis and found no significant differences in the amplitudes of scotopic a- and b-waves and photopic b-waves between *Mak^+/−^* and *Mak^+/−^*; *Ccrk^+/−^* mice, although the photopic a-wave amplitude significantly but slightly decreased in *Mak^+/−^*; *Ccrk^+/−^* mice compared with that in *Mak^+/−^* mice (Fig. S8B, C). These results show that *Ccrk* heterozygous deficiency alone hardly affects retinal photoreceptor function and maintenance.

Next, we performed histological analyses of retinal sections from *Mak^−/−^*; *Ccrk^+/−^* mice at 2M. Immunohistochemical examination showed more disorganized structures of rod outer segments in *Mak^−/−^*; *Ccrk^+/−^*mice than in *Mak^−/−^* mice (Fig. 6A). Mislocalization of cone opsins to the inner part of photoreceptors was more prominent in the *Mak^−/−^*; *Ccrk^+/−^* retina than in the *Mak^−/−^* retina (Fig. 6A). To evaluate the electrophysiological properties of *Mak^−/−^*; *Ccrk^+/−^* retina, we performed an ERG analysis and found that photopic a- and b-wave amplitudes in *Mak^−/−^*; *Ccrk^+/−^* mice were significantly lower than those in *Mak^−/−^* mice (Fig. 6B, C, and S8D, E). Although we observed that the photopic a-wave amplitudes to the brightest stimulus were decreased by *Ccrk* heterozygous deficiency in both *Mak^+/−^* and *Mak^−/−^* mice, the reduction rate was higher in *Mak^−/−^* mice (42%) than in *Mak^+/−^* mice (19%) (Fig. 6B, C, and S8B, C). These results suggest that *Ccrk* is a modifier of retinal degeneration observed in *Mak^−/−^* mice and support the idea that the Ccrk-Mak/Ick axis functions in retinal photoreceptor cells.

**Figure 6.**
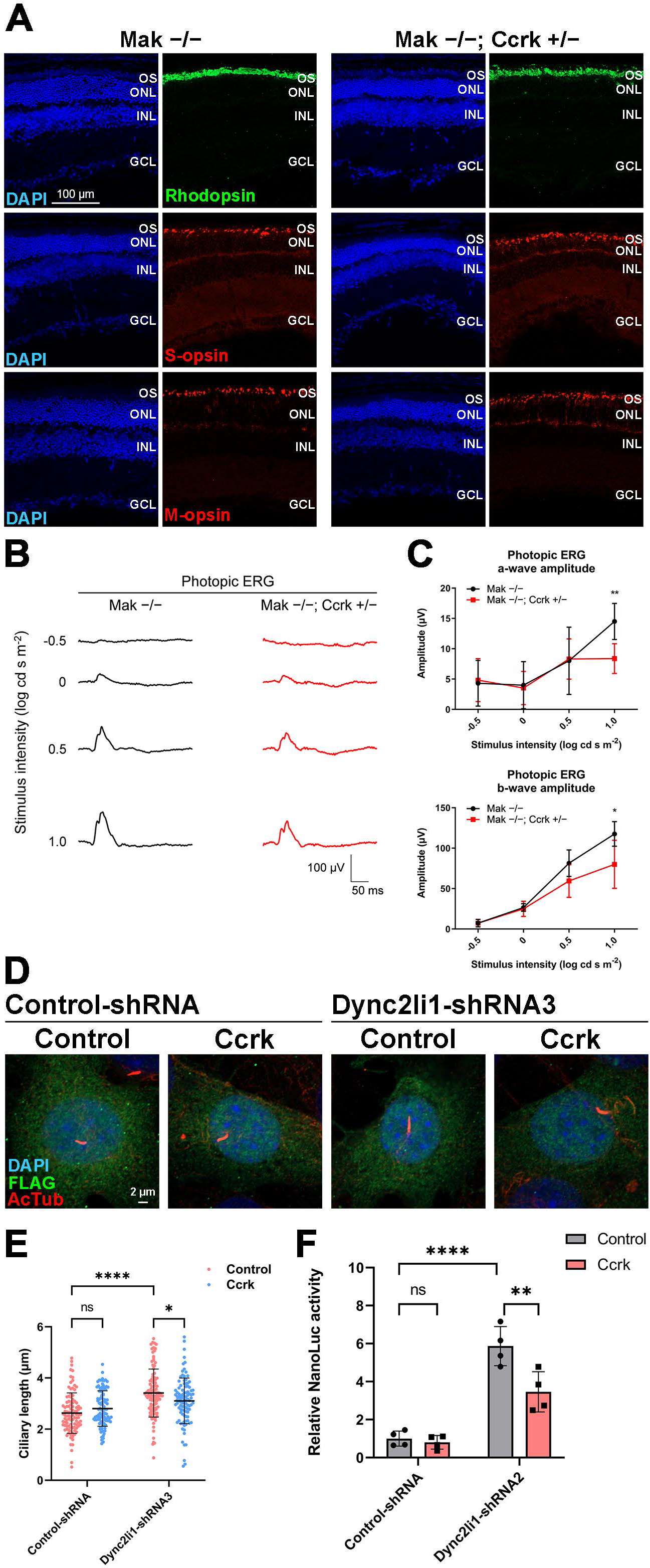
Effects of *Ccrk* dosage on retinal degeneration in *Mak^−/−^* mice and ciliary abnormalities caused by *Dync2li1* deficiency. (A) Immunostaining of retinal sections from *Mak^−/−^* and *Mak^−/−^*; *Ccrk^+/−^*mice at 2M using marker antibodies against Rhodopsin, S-opsin, and M-opsin. Rod outer segment disorganization and mislocalization of cone opsins were enhanced in the *Mak^−/−^*; *Ccrk^+/−^* retina. OS, outer segment; ONL, outer nuclear layer; INL, inner nuclear layer; GCL, ganglion cell layer. (B, C) ERG analysis of *Mak^−/−^*; *Ccrk^+/−^* mice at 2M. (B) Representative photopic ERGs elicited by four different stimulus intensities (−0.5 to 1.0 log cd s/m^2^) from *Mak^−/−^* and *Mak^−/−^*; *Ccrk^+/−^*mice. (C) The photopic amplitudes of a- and b-waves are shown as a function of the stimulus intensity. Data are presented as mean ± SD. \**p* < 0.05, \*\**p* < 0.01 (unpaired t-test). n = 7 and 4 mice (*Mak^−/−^* and *Mak^−/−^*; *Ccrk^+/−^*, respectively). (D, E) Effects of Ccrk overexpression on ciliary length in cells knocked down for *Dync2li1*. (D) A plasmid encoding Control-shRNA or Dync2li1-shRNA3 was co-transfected into NIH3T3 cells with or without a plasmid expressing Ccrk in combination with a construct encoding FLAG-tagged EGFP. Cells were immunostained with anti-FLAG and anti-Actub antibodies. (E) The length of cilia stained with an antibody against Actub in FLAG-positive cells was measured. Data are presented as mean ± SD. \**p* < 0.05, \*\*\*\**p* < 0.0001, ns, not significant (two-way ANOVA followed by Tukey’s multiple comparisons test). Control-shRNA; Control, Control-shRNA; Ccrk, Dync2li1-shRNA3; Control, and Dync2li1-shRNA3; Ccrk, n = 100, 104, 108, and 101 cilia, respectively, from 4 experiments. (F) Luciferase reporter gene assay using 8x Gli1-binding sites-minimal promoter-NanoLuc luciferase constructs. NIH3T3 cells were transfected with plasmids expressing Dync2li1-shRNA2 and Ccrk along with a Nanoluc luciferase reporter construct driven by the 8x Gli1-binding sites and minimal promoter and a Firefly luciferase-expressing construct driven by the SV40 promoter and enhancer. Luciferase activities of cell lysates were measured 24 h after serum starvation with 100 nM SAG. Nanoluc luciferase activity was normalized to Firefly luciferase activity. Data are presented as mean ± SD. \*\**p* < 0.01, \*\*\*\**p* < 0.0001, ns, not significant (two-way ANOVA followed by Tukey’s multiple comparisons test). N = 4 experiments. Nuclei were stained with DAPI.

To test whether Ccrk activation can restore the ciliary abnormalities caused by cytoplasmic dynein defects, we transfected the Control-shRNA or Dync2li1-shRNA3 expression plasmids into NIH3T3 cells with plasmids encoding Ccrk and examined the cilia by immunostaining using the anti-Actub antibody. We found that Ccrk expression rescued Dync2li1-shRNA-induced cilia elongation (Fig. 6D, E). We also observed that Ccrk expression suppresses *Dync2li1* knockdown-induced Hedgehog signaling activation (Fig. 6F). These results suggest that activation of the Ccrk-Mak/Ick axis can rescue ciliary defects caused by cytoplasmic dynein inhibition.

## Discussion

In the current study, we performed molecular, cellular, histological, and electrophysiological analyses in combination with mouse genetics and identified Ccrk-Mak/Ick kinase signaling as an IFT regulator essential for retinal photoreceptor maintenance. In addition, our results shed light on pathological mechanisms underlying retinitis pigmentosa caused by mutations in the human *MAK* gene and suggest that activation of Ick is a potential therapeutic approach to treat this disease. We previously reported that Ick regulates the IFT turnaround step at the tip of the cilia (28). IFT-A, IFT-B, and BBSome components are accumulated at the ciliary tips in *Ick^−/−^*MEFs (28). Although Ick is recognized as a critical regulator of the IFT turnaround process (16, 27), there were no obvious differences in the distribution of IFT components in the photoreceptor ciliary axonemes between the control and *Ick* CKO retinas. This observation prompted us to screen other serine-threonine kinase(s) and Mak was identified as a candidate regulator of IFT turnaround step. We observed that Mak and Ick localized to the ciliary tips in cultured cells and to the distal region of photoreceptor ciliary axonemes in the retina. It was previously reported that Ick localization to the ciliary tip is mediated through anterograde trafficking by IFT-B (39). A recent study showed a critical role of IFT-A·Tubby-like protein (TULP) carriages in the ciliary tip localization of Ick (16). Using these mechanisms, Mak and Ick may be transported to the distal region of the ciliary axonemes in the retinal photoreceptor cells. We also observed that the overexpression of Mak and Ick similarly changed the ciliary localization of IFT components in cultured cells. IFT components were concentrated at the tips of the photoreceptor ciliary axonemes in the *Mak^−/−^* retina, which is similar to observations in *Ick^−/−^* MEFs (28). Phosphorylation of Kif3a, an Ick substrate (28), was lower in the *Mak^−/−^* retina than in the *Mak^+/+^* retina. In addition, the deletion of both *Mak* and *Ick* leads to severe retinal degeneration. Based on these findings, we concluded that Mak and Ick play a central role in IFT turnaround regulation in retinal photoreceptor cells, although we cannot exclude the possibility that kinase(s) other than Ick and Mak function as regulators of the IFT turnaround process and that Mak and Ick may have functions other than the regulation of the IFT turnaround process. We previously observed that both *Mak* and *Ick* are expressed in the brain, lung, spleen, ovary, and testis, in addition to the retina (64), suggesting that *Mak* and *Ick* genetically interact and play a role in IFT regulation not only in the retina but also in other tissues. Non-syndromic retinitis pigmentosa, but not syndromic ciliopathies, exhibited by individuals with *MAK* gene mutations (49, 50) may be due to genetic interactions between *MAK* and *ICK*.

We found that Ick Thr-157 phosphorylation is critical for its kinase activity, which is consistent with previous studies indicating its importance in Mak and Ick activity (63, 65, 66). The level of Thr-157 phosphorylation in the Mak protein was markedly decreased in the *Ccrk* CKO retina compared to that in the control retina. *Ccrk* CKO mice exhibited progressive retinal degeneration phenotypes, quite similar to those observed in *Mak Ick* DKO mice. In addition, *Mak^−/−^*; *Ccrk^+/−^* retina were more severely degenerated than *Mak^−/−^*retina. These observations strongly suggest that Ccrk acts as a major upstream regulator of Mak and Ick in retinal photoreceptor cells, which is supported by previous reports showing that Ccrk directly phosphorylates Ick and Mak at Thr-157 *in vitro* (63, 66). However, we cannot exclude the possibility that Mak and Ick activity is regulated by other mechanism(s). It was previously reported that protein phosphatase 5 (PP5) dephosphorylates Ick at Thr-157 (66), implying that Mak and Ick are negatively regulated by PP5 in retinal photoreceptor cells. Fgfrs interact with, phosphorylate, and inactivate Ick (54) and are expressed in retinal photoreceptor cells (67–73). We observed that retinal degeneration in *Mak^−/−^* mice was suppressed by BGJ398, an FDA-approved inhibitor of Fgfrs, whereas that in *Mak Ick* DKO mice could not be rescued by BGJ398. This observation suggests that the protective role of BGJ398 on the *Mak^−/−^* retina is mediated through Ick activation, although we cannot exclude the possibility that other pathway(s) are also involved in the effects of BGJ398 treatment. The increased phosphorylation level of Kif3a in the retina of *Mak^−/−^* mice treated with BGJ398 supports the idea that Ick is activated in the *Mak^−/−^* retina by pharmacological inhibition of Fgfrs. These observations suggest that fibroblast growth factor-Fgfr signaling negatively regulates Ick activity in retinal photoreceptor cells, although Ick activation in retinal cells other than photoreceptors may have contributed to the suppression of retinal degeneration in BGJ398-treated *Mak^−/−^* mice.

Photoreceptor ciliary axonemes in *Mak^−/−^* retinas were elongated, whereas those in *Mak Ick* DKO retinas were not observed. Accumulation of IFT components at ciliary tips in cultured cells and cochlear hair cells lacking *Ick* (28, 38, 39) as well as photoreceptor cells of the *Mak^−/−^* retina shows that retrograde IFT can be inhibited by disruption of *Ick* and *Mak*. *Dync2li1* knockdown and knockout caused elongation and severe shortening of cilia in RPE1 cells, respectively (74, 75), suggesting that ciliary length is affected by the extent to which retrograde IFT is inhibited. In contrast to the absence or shortening of cilia caused by the loss of *Dync2li1*, a small amount of Dync2li1 protein remaining in the knockdown cells may induce cilia elongation. This idea is supported by ciliary elongation in cultured cells knocked down for *Dync2li1* in our study, and the absence of cilia or ciliary shortening in the ventral node of mouse embryos lacking *Dync2li1* in a previous study (76). Therefore, the differences in photoreceptor ciliary axonemes between the *Mak^−/−^*and *Mak Ick* DKO retinas may be attributed to the severity of retrograde IFT defects. Ick may compensate for the impairment of retrograde IFT in the *Mak^−/−^* retina. We and others have previously reported that loss of function of *Ick* results in elongation or shortening/absence of cilia (28, 36–39, 42, 43, 54), raising the possibility that the total remaining amount and/or activity of Ick and Mak in cells with elongated cilia is higher than that in cells showing shortening or absence of cilia. Previous studies have shown that IFT components accumulate at the ciliary tips in *Ccrk^−/−^* MEFs and *Ccrk*-deficient RPE1 cells (77, 78), supporting the idea that the Ccrk-Mak/Ick signaling pathway regulates the IFT turnaround process in retinal photoreceptor cells. Notably, cilia were elongated in *Ccrk^−/−^* MEFs and *Ccrk*-deficient RPE1 cells (77, 78), whereas photoreceptor ciliary axonemes were not observed in the *Ccrk* CKO retina. A possible explanation for this disparity is the differences between cultured cells and *in vivo* conditions or the severity of retrograde IFT defects. Dysfunction of retrograde IFT in *Ccrk^−/−^* MEFs and *Ccrk*-deficient RPE1 cells may be compensated in cultured conditions. Loss of function of *Ccrk* orthologues, *lf2* in *Chlamydomonas* and *dyf-18* in *C. elegans*, also causes flagella/cilia elongation (33, 34, 79, 80). Thus, in this case, it is unclear whether the data obtained from invertebrates and cultured cell systems can directly apply to mammals *in vivo*.

How do Mak and Ick regulate the IFT turnaround process at ciliary tips through phosphorylation in retinal photoreceptor cells? We have previously reported that Ick phosphorylates the C-terminal portion of Kif3a, including Thr-674 (28). We observed that Kif3a phosphorylation markedly decreased in the *Mak^−/−^* retina compared to that in the *Mak^+/+^*retina, suggesting that Mak phosphorylates Kif3a in retinal photoreceptor cells. Phosphorylation of serine/threonine residues, including Thr-674 in the Kif3a C-terminal region, is required for cilia formation in cultured cells and zebrafish (28). In contrast, MEFs harboring a Thr-to-Ala mutation at residue 674 on Kif3a exhibited slight cilia elongation without affecting the ciliary localization of IFT88 (81). In *Chlamydomonas*, inhibition of phosphorylation of the kinesin-2 motor subunit FLA8, an ortholog of Kif3b, at Ser-663 led to defects in IFT turnaround at the flagellar tip (82). FLA8 Ser-663 is located in a consensus amino acid sequence for phosphorylation by Ick (66) which is evolutionarily conserved among species, implying that Kif3b phosphorylation by Mak and Ick modulates IFT turnaround at the ciliary tip in retinal photoreceptor cells. *C. elegans* DYF-5 reduced the tubulin-binding affinity of the IFT-B components IFT-74/81 by phosphorylating IFT74, proposing a model in which DYF-5-mediated phosphorylation of IFT74 promotes tubulin unloading at the ciliary tip (83–85). Future studies are needed to uncover the downstream mechanisms underlying the regulation of IFT by Mak and Ick in retinal photoreceptor cells.

Some of retinitis pigmentosa patients with the same mutation in the *MAK* gene represent different severities of disease (86). We found that *Mak^−/−^*; *Ick^+/−^* mice exhibited severe retinal degeneration compared with *Mak^−/−^*mice, and that *MAK* and *ICK* are expressed in photoreceptor cells of the human retina. Heterozygous loss-of-function mutations/variants in the human *ICK* gene (41–44) may cause more severe symptoms in retinitis pigmentosa patients having *MAK* mutations. Additionally, mutations or variants in other genes involved in retrograde IFT may contribute to the severity of retinal degeneration caused by *MAK* mutations. Indeed, we observed that *Mak^−/−^*; *Ccrk^+/−^* retinas were more severely degenerated than *Mak^−/−^* retinas. Ccrk physically and functionally interacts with Bromi (60). We found that both *CCRK* and *BROMI* were expressed in photoreceptor cells of the human retina. Notably, mutations in the human *BROMI* gene are associated with progressive retinal dystrophy (87). Our results may advance our understanding of the pathological mechanisms underlying progressive retinal dystrophy in patients with mutations in the human *BROMI* gene.

We found that Mak can rescue ciliary abnormalities caused by *Ick* deficiency and *vice versa*. In addition, BGJ398 and the overexpression of Ick, Mak, and Ccrk could rescue ciliary defects due to the deficiency of *Dync2li1*, a ciliopathy gene encoding a subunit of cytoplasmic dynein 2 (75), which supports the idea that the Ccrk-Mak/Ick axis engages in the turnaround process and subsequent retrograde IFT. These results suggest that ciliopathies defective in these IFT processes can be restored by the activation of the CCRK-MAK/ICK signaling pathway. Although the precise functional relationship between the Ccrk-Mak/Ick pathway and IFT-A/cytoplasmic dynein-2 remains unclear, mutations in the genes encoding IFT-A and cytoplasmic dynein-2, and *ICK* are associated with a shared human ciliopathy, SRPS (88). Further mechanistic studies would reveal the extent to which our proposed therapeutic approach can be more generally applied to treat ciliopathies.

## Materials and methods

### Animal care

All procedures conformed to the ARVO Statement for the Use of Animals in Ophthalmic and Vision Research, and were approved by the Institutional Safety Committee on Recombinant DNA Experiments (approval ID 04913) and the Animal Experimental Committees of the Institute for Protein Research (approval ID R04-02-0), Osaka University, and were performed in compliance with the institutional guidelines. Mice were housed in a temperature-controlled room at 22°C with a 12 hr light/dark cycle. Fresh water and rodent diet were available at all times. All animal experiments were performed with mice of either sex.

### Mouse lines

To generate the *Ccrk* flox mouse line, we subcloned an ∼ 11.5 kb *Ccrk* genomic fragment using C57BL/6 genomic DNA by PCR, inserted one loxP site into intron 2 and another loxP site into intron 4, cloned it into a modified pBluescript II KS (+) vector (Agilent) to make a targeting construct, and transfected the linearized targeting construct into the JM8A3 embryonic stem (ES) cell line (89). Culture, electroporation, and selection of JM8A3 cells were performed as described previously (61). ES cells heterozygous for targeted gene disruption were microinjected into C57BL/6 blastocysts to obtain chimeric mice. These chimeric mice were bred with C57BL/6 mice to obtain their progenies, which were subsequently crossed with B6-Tg(CAG-FLPe)37 mice (#RBRC01835, RIKEN BRC) to remove the flippase recognition target (FRT)-flanked neo cassette using flippase (Flp) recombinase. We obtained *Ccrk^+/−^* mice by crossing the *Ccrk* flox mouse line with the female *BAC-Dkk3-Cre* transgenic mouse line, whose progenies exhibit complete recombination in all tissues, irrespective of the co-transmission of the *BAC-Dkk3-Cre* transgene (45). Primer sequences used for genotyping are listed in Table S1. *Ick* flox mice (28), *Ick^−/−^* mice (28), *Cdkl5^−/Y^* mice (90), *Mak^−/−^* mice (91), and *Crx-CreERT2* transgenic mice (61) have been described previously.

### Tamoxifen treatment

Tamoxifen (Toronto Research Chemicals) was dissolved in sunflower seed oil (Sigma) to a concentration of 20 mg/ml and injected 4 mg of it intraperitoneally into mice at 1M. The injected mice were analyzed at 2M.

### RT-PCR analysis

RT-PCR analysis was performed as described previously (92). Total RNAs were extracted using TRIzol (Ambion) from tissues dissected from ICR mice at 4wks, retinas from the control and *Ccrk* CKO mice at P14, and retinas from the control and *Ccrk* iCKO mice at 2M. Total RNA (2 μg) was reverse-transcribed into cDNA with random hexamers using the PrimeScript II reagent (TaKaRa). Human cDNA was purchased from Clontech. The cDNAs were used for PCR with rTaq polymerase (TaKaRa). The primer sequences used for amplification are listed in Table S1.

### *In situ* hybridization

*In situ* hybridization was performed as described previously (93). Mouse eye cups were fixed using 4% paraformaldehyde (PFA) in phosphate-buffered saline (PBS) for 30 min at room temperature. The tissues were equilibrated in 30% sucrose in PBS overnight at 4°C, embedded in TissueTec OCT compound 4583 (Sakura, Japan), and frozen. Digoxigenin-labeled antisense riboprobes for mouse *Ccrk* were synthesized by *in vitro* transcription using 11-digoxigenin UTPs (Roche). A *Ccrk* cDNA fragment for *in situ* hybridization probe was generated by PCR, using mouse testicular cDNA as a template. The primer sequences used for amplification are listed in Table S1.

### Plasmid constructs

Plasmids expressing EGFP, FLAG- or HA-tagged mouse Mak and Ick, FLAG-tagged EGFP, mouse IFT57, IFT88, IFT140, BBS8, and Ick-KD, and HA-tagged human ICK were previously constructed (28, 47, 94). Full-length cDNA fragments of mouse *Cdkl1*, *Cdkl2*, *Cdkl3*, *Cdkl4*, *Cdkl5*, *Gsk3b*, and *Dync2li1* were amplified by PCR using mouse retinal cDNA as a template and subcloned into the pCAGGSII-3xFLAG and/or pCAGGSII-2xHA (62, 95) vectors. Full-length cDNA fragments of mouse *Mok*, *Gsk3a* and *Ccrk* were amplified by PCR using mouse testicular cDNA as a template and subcloned into the pCAGGSII-3xFLAG and/or pCAGGSII (47) vectors. Full-length cDNA fragment of human *MAK* was amplified by PCR using human retinal cDNA (Clontech) as a template and subcloned into the pCAGGSII-2xHA vector (62). The mouse Ick T157A mutation was introduced via site-directed mutagenesis using PCR. For shRNA-mediated *Dync2li1* knockdown, Dync2li1-shRNA and Control-shRNA cassettes were subcloned into the pBAsi-mU6 vector (Takara Bio, Shiga, Japan). The target sequences were as follows: Dync2li1-shRNA2, 5′-GCTTTGTGGCACATTACTACG-3′; Dync2li1-shRNA3, 5′-GCAGGACTGGATTCTTTATGT-3′; Dync2li1-shRNA5, 5′-GGGAATTAATTGACCCATTTC-3′; Dync2li1-shRNA6, 5′-GCAAGTCAGAAGCTCTGTTAC-3′; Control-shRNA, 5′-GACGTCTAACGGATTCGAGCT-3′ (96). To produce AAV-Rhodopsin kinase promoter-3xFLAG-Ick, a full-length cDNA fragment of mouse *Ick* was amplified by PCR using pCAGGSII-2xHA-mouse Ick as a template and subcloned into the pBluescript II KS (+)-3xFLAG vector, which was modified from the pBluescript II KS (+) vector. *3xFLAG-Ick* digested from pBluescript II KS (+)-3xFLAG-Ick was ligated into the pAAV-RK-IZsGreen vector (a kind gift from Dr. T. Li) (53). The primer sequences used for the amplification are listed in Table S1.

### Chemicals

BGJ398 was purchased from ChemScene. Smoothened agonist (SAG) was obtained from Calbiochem. Ciliobrevin D was purchased from Millipore.

### Cell culture and transfection

HEK293T and NIH3T3 cells were cultured in Dulbecco’s modified Eagle’s medium (DMEM) (Sigma) containing 10% fetal bovine serum and calf serum, respectively, supplemented with penicillin (100 μg/ml) and streptomycin (100 μg/ml), at 37°C with 5% CO_2_. *Ick^−/−^* MEFs were derived from embryonic day 13.5 embryos and cultured in DMEM containing 10% fetal bovine serum supplemented with penicillin (100 μg/ml) and streptomycin (100 μg/ml) at 37°C with 5% CO_2_. Transfection was carried out using the calcium phosphate method for HEK293T cells and Lipofectamine LTX (Invitrogen) or Lipofectamine 3000 (Invitrogen) for NIH3T3 cells and *Ick^−/−^* MEFs. To induce ciliogenesis in transfected cells, the medium was replaced with serum-free medium 24 h after transfection, and the cells were cultured for 24 h in serum-free medium. NIH3T3 cells were incubated with 100 nM BGJ398, 100 nM SAG, or 10 μM Ciliobrevin D for 24 h in serum-free medium.

### Immunofluorescent analysis of cells and retinal sections

Immunofluorescence analysis of the cells and retinal sections was performed as described previously (97, 98). Cells were washed with PBS, fixed with 4% PFA in PBS for 5 min at room temperature, and subsequently incubated with blocking buffer (5% normal donkey serum and 0.1% or 0.5% Triton X-100 in PBS) for 30 min at room temperature, and then immunostained with primary antibodies in blocking buffer overnight at 4°C. Cells were washed with PBS and incubated with secondary antibodies and DAPI (1:1,000, Nacalai Tesque) in blocking buffer for 2 h at room temperature. After washing three times with PBS, the samples were coverslipped with gelvatol. The mouse eyes or eye cups were fixed with 4% PFA in PBS for 15 sec to 30 min at room temperature. Eye samples were rinsed in PBS, embedded in TissueTec OCT compound 4583 (Sakura), frozen, and sectioned. The eye cup samples were rinsed in PBS, followed by cryoprotection using 30% sucrose in PBS overnight at 4°C, embedded in TissueTec OCT compound 4583, frozen, and sectioned. Frozen 14 μm or 20 μm sections on slides were dried overnight at room temperature, rehydrated in PBS for 5 min, incubated with blocking buffer for 30 min, and then with primary antibodies overnight at 4°C. Slides were washed with PBS three times for 5 min each and incubated with fluorescent dye-conjugated secondary antibodies and DAPI for 2 h at room temperature while shielded from light. After washing three times with PBS, the sections were coverslipped with gelvatol. The specimens were observed under a laser confocal microscope (LSM700, LSM710, or LSM900; Carl Zeiss). Primary antibodies used in this study were as follows: mouse anti-Actub (1:1,000 or 1:2,000, Sigma, 6-11B-1), guinea pig anti-Ick (1:100) (28), rabbit anti-IFT88 (1:500, Proteintech, 13967-1-AP), rabbit anti-IFT140 (1:500, a kind gift from Dr. G. J. Pazour) (99), rabbit anti-FLAG (1:1,000, Sigma, F7425), mouse anti-FLAG-M2 (Sigma, F1804, 1:1,000), mouse anti-γ-tubulin (1:250, Sigma, T6557), rabbit anti-pericentrin (1:1,000, Abcam, ab4448), guinea pig anti-Mak (1:1,000) (47), mouse anti-Cep164 (1:200, Santa Cruz, sc-515403), rabbit anti-Rhodopsin (1:2,500, LSL, LB-5597), goat anti-S-opsin (1:500, Santa Cruz, sc-14363 or 1:100, ROCKLAND, 600-101-MP7S), rabbit anti-M-opsin (1:500, Millipore, AB5405), and rabbit anti-(pThr^674^) Kif3a (1:250) (28). Cy3-conjugated (1:500, Jackson ImmunoResearch Laboratories) and Alexa Fluor 488-conjugated (1:500, Sigma) secondary antibodies were used. Ciliary length was measured using ZEN imaging software (ZEN (blue edition), Carl Zeiss). The rod outer segment lengths and immunofluorescence signal intensities were measured and quantified using National Institutes of Health (NIH) ImageJ software. The Rhodopsin signals in the inner part of the photoreceptors were normalized to the total (OS + inner part) signals for Rhodopsin.

### Toluidine blue staining

Toluidine blue staining of the retinal sections was performed as described previously (100). The retinal sections were rinsed with PBS and stained with 0.1% toluidine blue in PBS for 1 min. After washing with PBS, the slides were covered with coverslips and immediately observed under a microscope. Retinal thickness was measured and quantified using NIH ImageJ software as described previously (94).

### Transmission electron microscope analysis

Transmission electron microscopy (TEM) was performed as previously described (101). Mouse eye cups were fixed with 2% glutaraldehyde and 2% PFA in PBS for 30 min. Retinas were fixed with 1% osmium tetroxide for 90 min, dehydrated through a graded series of ethanol (50%–100%), cleared with propylene oxide, and embedded in epoxy resin. Sections were cut on an ultramicrotome (Ultracut E; Reichert-Jung), stained with 2% uranyl acetate and Sato’s lead staining solution (102), and observed under a transmission electron microscope (H-7500; Hitachi Co).

### ERG recording

Electroretinograms (ERGs) were recorded as described previously (103). Briefly, mice were dark-adapted overnight and then anesthetized with an intraperitoneal injection of 100 mg/kg ketamine and 10 mg/kg xylazine diluted in saline (Otsuka). The pupils were dilated using topical 0.5% tropicamide and 0.5% phenylephrine HCl. The ERG responses were measured using a PuREC system with LED electrodes (Mayo Corporation). Mice were placed on a heating pad and stimulated with an LED flash. Four levels of stimulus intensities ranging from −4.0 to 1.0 log cd s m^−2^ were used for the scotopic ERGs. After the mice were light-adapted for 10 min, the photopic ERGs were recorded on a rod-suppressing white background of 1.5 log cd m^−2^. Four levels of stimulus intensities ranging from −0.5 to 1.0 log cd s m^−2^ were used for the photopic ERGs. Eight responses at −4.0 log cd s m^−2^ and four responses at −3.0 log cd s m^−2^ were averaged for the scotopic recordings. Sixteen responses were averaged for the photopic recordings.

### Western blot analysis

Western blot analysis was performed as described previously (104). HEK293T cells were washed twice with Tris-buffered saline (TBS) and lysed in SDS-Sample buffer or a lysis buffer supplemented with protease inhibitors (Buffer A:20 mM Tris-HCl pH 7.4, 150 mM NaCl, 1% NP-40, 1 mM EDTA, 1 mM PMSF, 2 µg/ml leupeptin, 5 µg/ml aprotinin, and 3 µg/ml pepstatin A). Mouse retinas were lysed in buffer A or B (20 mM Tris-HCl pH 7.4, 150 mM NaCl, and 1% NP-40 supplemented with phosphatase inhibitor cocktail (Roche)). Samples were resolved by SDS-PAGE and transferred to PVDF membranes (Millipore) using a semidry transfer cell (Bio-Rad) or iBlot system (Invitrogen). The membranes were blocked with blocking buffer (3% skim milk or 1% bovine serum albumin, and 0.05% Tween 20 in TBS) and incubated with primary antibodies overnight at 4°C. The membranes were washed with 0.05% Tween 20 in TBS three times for 10 min each and then incubated with secondary antibodies for 2 h at room temperature. The signals were detected using Chemi-Lumi One L (Nacalai) or Pierce Western Blotting Substrate Plus (Thermo Fisher Scientific). The following primary antibodies were used: rabbit anti-Kif3a (1:1,500, Abcam, ab11259), mouse anti-FLAG M2 (1:5,000 or 1:10,000, Sigma, F1804), rabbit anti-GFP (1:2,500, MBL, 598), rabbit anti-pMak (1:1,000, Invitrogen, PA5-105526), guinea pig anti-Mak (1:150) (47), guinea pig anti-Ccrk (1:500, generated in this study), and mouse anti-α-tubulin (Cell Signaling, DM1A, 1:5,000, T9026). The following secondary antibodies were used: horseradish peroxidase-conjugated anti-mouse IgG (1:10,000; Zymed), anti-guinea pig IgG (1:10,000; Jackson Laboratory), and anti-rabbit IgG (1:10,000; Jackson Laboratory). Phosphorylated Kif3a was detected by 6% SDS-PAGE with 20 µM Phos-tag acrylamide (FUJIFILM Wako), according to the manufacturer’s instruction (105, 106). The band intensity was quantified using NIH ImageJ software.

### Antibody production

Antibody production was performed as described previously (107). Briefly, a cDNA fragment encoding the C-terminal portion of mouse Ccrk (residues 289-346) was amplified by PCR and subcloned into the pGEX-4T-3 vector (GE Healthcare). The GST-fused Ccrk protein was expressed in *Escherichia coli* strain BL21 (DE3) and purified using glutathione Sepharose 4 B (GE Healthcare). An antibody against Ccrk was obtained by immunizing guinea pigs with purified GST-fused Ccrk.

### Analysis of scRNA-seq data

To analyze the publicly available scRNA-seq data of the adult human retina (52), a processed data (ae_exp_proc_all.tsv) obtained from http://www.ebi.ac.uk/arrayexpress/experiments/E-MTAB-7316 was imported into the R package Seurat (version 4.3.0) (108). The data were normalized using the LogNormalize method with a scale factor of 10,000, and the variable features were identified using FindVariableFeatures with 2,000 genes. Dimensionality reduction was then performed using PCA. Cell clustering was performed using FindNeighbors and FindClusters (dim = 20, resolution = 0.6). UMAP was performed to visualize cell clusters using the RunUMAP function. Cell identities were assigned to each cluster as described previously (52).

### AAV production and subretinal injection

AAV production and subretinal injection of AAVs were performed as previously described (109). AAVs were produced by triple transfection of an AAV vector plasmid, adenovirus helper plasmid, and AAV helper plasmid (pAAV2-8 SEED) into AAV-293 cells using the calcium phosphate method. The cells were harvested 72 h after transfection and lysed using four freeze-thaw cycles. The supernatant was collected by centrifugation and treated with Benzonase nuclease (Novagen). The viruses were purified using iodixanol gradient. The gradient was formed in ultra-clear centrifuge tubes (14×95 mm, Beckman) by first adding 54% iodixanol (Axis-Shield) in PBS-MK buffer (1×PBS, 1 mM MgCl_2_, and 25 mM KCl), followed by overlaying 40% iodixanol in PBS-MK buffer, 25% iodixanol in PBS-MK buffer containing phenol red, and 15% iodixanol in PBS-MK buffer containing 1 M NaCl. The tubes were centrifuged at 40,000 rpm using an SW40Ti rotor. The 54%-40% fraction containing the virus was collected using an 18-gauge needle and concentrated using an Amicon Ultra Centrifugal Filter Ultracel-100K (Millipore). The titer of each AAV (in vector genomes (VG)/ml) was determined by qPCR using SYBR GreenER Q-PCR Super Mix (Invitrogen) and Thermal Cycler Dice Real Time System Single MRQ TP700 (Takara) according to the manufacturer’s protocols. Quantification was performed using Thermal Cycler Dice Real Time System software (Takara). The primers used for AAV titrations are listed in Table S1. The titer of AAV-RK-N-FLAGx3-Ick was adjusted to approximately 1.8×10^12^ VG/ml. 0.3 µl of the AAV preparation was injected into the subretinal spaces of P1 *Mak^−/−^* mice. The injected retinas were harvested at P14.

### Drug administration

BGJ398 dissolved in DMSO (2 or 4 mg/ml) was diluted in 3.5 mM HCl and 5% DMSO. Two mg/kg of BGJ398 was administered to mice every day by subcutaneous injection.

### Luciferase reporter assay

Reporter gene assays were performed using the Nano-Glo Dual-Luciferase Reporter Assay System (Promega) according to the manufacturer’s protocol. To generate the reporter construct, a minimal promoter (5′-AGACACTAGAGGGTATATAATGGAAGCTCGACTTCCAG-3′) and 8x Gli1-binding sites (110) were cloned into the pGL3-Basic vector (Promega), in which *Firefly luciferase* was replaced with *NanoLuc luciferase* amplified from the pNLF-N [CMV/Hygro] vector (Promega) by PCR. The pGL3-Control vector (Promega) was co-transfected to normalize the transfection efficiency. Plasmids expressing Dync2li1-shRNA2, Dync2li1-shRNA3, or Dync2li1-shRNA6, and Ick, Mak iso1, Mak iso2, or Ccrk were transfected with the reporter constructs into NIH3T3 cells. After 24 h of transfection, cells were incubated in serum-free medium containing 100 nM SAG for 24 h and washed with PBS. Luminescence signals were detected using the GloMax® Multi+ Detection System (Promega).

### Sequence alignment and structure modelling

Structure-based sequence alignment of mouse Mak and Ick was performed using Clustal Omega (111) and ESPript (112). Mouse Mak and Ick 3D structures were obtained from the AlphaFold Protein Structure Database developed by DeepMind, in which the protein structure was predicted based on the amino acid sequence (https://alphafold.ebi.ac.uk/) (113, 114). The predicted 3D structures of the kinase domains (residues 4-284) in Mak and Ick proteins were visualized and aligned using PyMOL.

### Statistical analysis

Data are represented as the mean ± SD. Statistical analysis was performed using unpaired *t*-test, one-way ANOVA, or two-way ANOVA, as indicated in the figure legends. A value of *p* < 0.05 was taken to be statistically significant.

## Acknowledgements

We thank Y. Shinkai for the *Mak^‒/‒^* mouse, G. J. Pazour for the anti-IFT140 antibody, T. Li for the pAAV-RK-IZsGreen vector, and M. Kadowaki, A. Tani, A. Ishimaru, T. Tsujii, M. Wakabayashi, Y. Nakamura, Y. Kinooka, K. Yoshida, S. Okochi, K. Fukunaga, S. Kubo, H. Yamamoto, Y. Sugita, L. R. Varner, R. Saito, K. Kobayashi, T. Yasui, M. Nishi, K. Wada, T. Takeda, R. Shimizu, S. Zhou, and K. Nakamura for technical assistance. This work was supported by Grant-in-Aid for Scientific Research (21H02657, 24K09996) and Grant-in-Aid for Challenging Research (Exploratory) (23K18199) from the Japan Society for the Promotion of Science, AMED-CREST (21gm1510006) from the Japan Agency for Medical Research and Development, Japan Science and Technology Agency (JST) Moonshot R&D (JPMJMS2024), JST COI-NEXT (JPMJPF2018), The Takeda Science Foundation, and The Uehara Memorial Foundation.

## Author contributions

TC and TF designed the project. TF generated *Ccrk* flox mice. TT generated *Cdkl5^−/Y^* mice. TC, YM, RT, MA, YM, and MN performed molecular biological and cell culture experiments. TC, YM, RT, MA, YM, MN, and NK carried out histological experiments. TC, YM, and RT performed ERG experiments. TC and YM carried out AAV infection experiments. TC, YM, and TF wrote the manuscript. TF supervised the project.

