## Supplemental information for "Ccrk-Mak/Ick kinase signaling axis is a ciliary transport regulator essential for retinal photoreceptor maintenance"

#### Supplemental figure legends

**Figure S1. Phenotypic analysis of the retina from *Ick* CKO mice at 1M.**

(A) Immunostaining of retinal sections from the control and *Ick* CKO mice using anti-Ick and anti-acetylated α-tubulin (Actub) (a marker for the ciliary axoneme) antibodies. Ick signals were detected at the distal region of ciliary axonemes in photoreceptor cells of the control retina but not of the *Ick* CKO retina.

(B) Ciliary localization of IFT components in photoreceptor cells of the *Ick* CKO retina. Retinal sections obtained from the control and *Ick* CKO mice were immunostained using antibodies against IFT88 (an IFT-B component), IFT140 (an IFT-A component), and Actub.

(C) Immunostaining of retinal sections from the control and *Ick* CKO mice using marker antibodies against Rhodopsin (rod outer segments), S-opsin (S-cone outer segments), and M-opsin (M-cone outer segments).

(D, E) ERG analysis of *Ick* CKO mice. (D) Representative scotopic and photopic ERGs elicited by four different stimulus intensities (–4.0 to 1.0 log cd s/$m^2$ and –0.5 to 1.0 log cd s/$m^2$, respectively) from the control and *Ick* CKO mice. (E) The scotopic and photopic amplitudes of a- and b-waves are shown as a function of the stimulus intensity. Data are presented as mean ± SD. The amplitudes of a- and b-waves were not significantly different between the control and *Ick* CKO mice (unpaired t-test). n = 6 and 4 mice (control and *Ick* CKO, respectively).

Nuclei were stained with DAPI. OS, outer segment; IS, inner segment; ONL, outer nuclear layer; INL, inner nuclear layer; GCL, ganglion cell layer.

**Figure S2. Phenotypic analysis of the retina from *Ick* CKO mice at 6M.**

(A-C) Measurement of retinal thickness in the control and *Ick* CKO mice at P14, 1M, and 6M. (A) Toluidine blue staining of retinal sections from the control and *Ick* CKO mice at P14, 1M, and 6M. (B) The ONL thickness was measured. Data are presented as mean ± SD. *$p < 0.05$, **$p < 0.01$, ***$p < 0.001$ (unpaired t-test). n = 4 and 5 mice at P14, n = 6 and 4 mice at 1M (control and *Ick* CKO, respectively), and n = 4 mice per each genotype at 6M. (C) Relative ONL thickness was expressed as a proportion of the ONL thickness to the INL+IPL+GCL thickness. Data are presented as mean ± SD. ***$p < 0.001$ (unpaired t-test). n = 4 and 5 mice at P14, n = 6 and 4 mice at 1M (control and *Ick* CKO, respectively), and n = 4 mice per each genotype at 6M.

(D) Immunostaining of retinal sections from the control and *Ick* CKO mice using marker antibodies against Rhodopsin, S-opsin, and M-opsin. Nuclei were stained with DAPI. Arrowheads indicate mislocalization of M-opsin to the OPL in the *Ick* CKO retina.

(E, F) ERG analysis of *Ick* CKO mice. (E) Representative scotopic and photopic ERGs elicited by four different stimulus intensities (–4.0 to 1.0 log cd s/$m^2$ and –0.5 to 1.0 log cd s/$m^2$, respectively) from the control and *Ick* CKO mice. (F) The scotopic and photopic amplitudes of a- and b-waves are shown as a function of the stimulus intensity. Data are presented as mean ± SD. *$p < 0.05$, **$p < 0.01$, ***$p < 0.001$ (unpaired t-test). n = 4 mice per each genotype.

OS, outer segment; ONL, outer nuclear layer; OPL, outer plexiform layer; INL, inner nuclear layer; IPL, inner plexiform layer; GCL, ganglion cell layer.

**Figure S3. Effects of Cdkl5 and Mak overexpression on ciliary localization of IFT components, and ERG analysis of *Cdkl5*$^{-/Y}$ mice.**

(A) Effects of Cdkl5, Mak, and Ick overexpression on ciliary localization of IFT57. A FLAG-tagged IFT57 expression plasmid was co-transfected into NIH3T3 cells with a plasmid expressing Cdkl5, Mak, or Ick. Cells were immunostained with anti-FLAG and anti-Actub antibodies. Arrowheads indicate ciliary tips. IFT57 localization to the ciliary tips increased by Mak or Ick but not Cdkl5 overexpression.

(B) RT-PCR analysis of the *Cdkl5* transcript in mouse tissues at 4wks. *β-actin* was used as a loading control.

(C) ERG analysis of *Cdkl5*$^{-/Y}$ mice. Representative scotopic and photopic ERGs elicited by four different stimulus intensities (–4.0 to 1.0 log cd s/m$^2$ and –0.5 to 1.0 log cd s/m$^2$, respectively) from *Cdkl5*$^{+/Y}$ and *Cdkl5*$^{-/Y}$ mice at 1M and 3M are shown. There were no substantial differences between *Cdkl5*$^{+/Y}$ and *Cdkl5*$^{-/Y}$ mice.

(D) Effects of Mak and Ick overexpression on ciliary localization of IFT140. A FLAG-tagged EGFP or IFT140 expression plasmid was co-transfected into NIH3T3 cells with a plasmid expressing Mak or Ick. Arrowheads indicate ciliary tips. IFT140 localization to the ciliary base increased by Mak or Ick overexpression.

(E) Effects of Mak and Ick overexpression on ciliary localization of BBS8. A FLAG-tagged EGFP or BBS8 expression plasmid was co-transfected into NIH3T3 cells with a plasmid expressing Mak or Ick. Arrowheads indicate ciliary tips. There were no substantial differences in the ciliary localization of BBS8 among control, Mak-, and Ick-overexpressing cells.

(F, G) Effects of human MAK and ICK overexpression on ciliary localization of IFT57. (F) A FLAG-tagged EGFP or IFT57 expression plasmid was co-transfected into NIH3T3 cells with a plasmid expressing human MAK or ICK. Cells were immunostained with anti-FLAG and anti-Actub antibodies. Arrowheads indicate ciliary tips. (G) The number of cilia with the FLAG-tagged EGFP or IFT57 predominantly localized at ciliary tips was counted. Data are presented as mean ± SD. ****$p < 0.0001$ (two-way ANOVA followed by Tukey's multiple comparisons test). n = 3 experiments.

Nuclei were stained with DAPI.

**Figure S4. Histological analysis of the *Mak Ick* DKO mouse retina.**

(A) Immunostaining of retinal sections from the control, $Mak^{-/-}$, and *Mak Ick* DKO mice at 1M using marker antibodies against Rhodopsin, S-opsin, and M-opsin. Severe photoreceptor degeneration was observed in the *Mak Ick* DKO retina.

(B) Cilia formation in photoreceptor cells of the *Mak Ick* DKO retina. Retinal sections obtained from the control, $Mak^{-/-}$, and *Mak Ick* DKO mice at P9 and P14 were immunostained using antibodies against Pericentrin (a marker for basal bodies) and Actub. Nuclei were stained with DAPI. OS, outer segment; IS, inner segment; ONL, outer nuclear layer; INL, inner nuclear layer; GCL, ganglion cell layer.

**Figure S5. Phenotypic analysis of the $Mak^{+/-}$; $Ick^{+/-}$ mouse retina.**

(A) RT-PCR analysis of the *MAK* and *ICK* transcripts in the human retina. *β-actin* was used as a loading control.

(B, C) Toluidine blue staining of retinal sections from $Mak^{+/+}$; $Ick^{+/+}$ and $Mak^{+/-}$; $Ick^{+/-}$ mice at 6M. Data are presented as mean ± SD. ns, not significant (unpaired t-test). n = 4 and 3 mice ($Mak^{+/+}$; $Ick^{+/+}$ and $Mak^{+/-}$; $Ick^{+/-}$, respectively).

(D) Immunostaining of retinal sections from $Mak^{+/+}$; $Ick^{+/+}$ and $Mak^{+/-}$; $Ick^{+/-}$ mice at 6M using marker antibodies against Rhodopsin, S-opsin, and M-opsin.

(E, F) ERG analysis of $Mak^{+/-}$; $Ick^{+/-}$ mice at 6M. (E) Representative scotopic and photopic ERGs elicited by four different stimulus intensities (−4.0 to 1.0 log cd s/m$^2$ and −0.5 to 1.0 log cd s/m$^2$, respectively) from *Mak*$^{+/+}$; *Ick*$^{+/+}$ and *Mak*$^{+/–}$; *Ick*$^{+/–}$ mice. (F) The scotopic and photopic amplitudes of a- and b-waves are shown as a function of the stimulus intensity. Data are presented as mean ± SD. The amplitudes of a- and b-waves were not significantly different between *Mak*$^{+/+}$; *Ick*$^{+/+}$ and *Mak*$^{+/–}$; *Ick*$^{+/–}$ mice (unpaired t-test). n = 4 and 3 mice (*Mak*$^{+/+}$; *Ick*$^{+/+}$ and *Mak*$^{+/–}$; *Ick*$^{+/–}$, respectively).

Nuclei were stained with DAPI. OS, outer segment; ONL, outer nuclear layer; INL, inner nuclear layer; GCL, ganglion cell layer.

**Figure S6. Retinal phenotypes in the mice treated with an Fgfr inhibitor, and effects of Ick overexpression on ciliary abnormities caused by cytoplasmic dynein inhibition.**

(A) Schematic diagram of schedule for drug administration. BGJ398 was injected into *Mak Ick* DKO and *Mak*$^{+/+}$ mice from P7 to 1M every day. s.c., subcutaneous.

(B, C) Toluidine blue staining of retinal sections from 1M *Mak Ick* DKO mice treated with or without BGJ398, which was injected into the mice from P7 to 1M every day. Data are presented as mean ± SD. ns, not significant (unpaired t-test). n = 5 and 6 mice (vehicle-treated and BGJ398-treated, respectively).

(D) ERG analysis of 1M *Mak Ick* DKO mice treated with BGJ398, which was injected into the mice from P7 to 1M every day. Representative scotopic ERGs elicited by two different stimulus intensities (−1.0 and 1.0 log cd s/m$^2$) from 1M *Mak Ick* DKO mice treated with or without BGJ398 are shown. There were no substantial differences between vehicle-treated and BGJ398-treated *Mak Ick* DKO mice.

(E, F) Toluidine blue staining of retinal sections from 1M *Mak*$^{+/+}$ mice treated with or without BGJ398, which was injected into the mice from P7 to 1M every day. Data are presented as mean ± SD. ns, not significant (unpaired t-test). n = 5 and 6 mice (vehicle-treated and BGJ398-treated, respectively).

(G) ERG analysis of 1M *Mak*$^{+/+}$ mice treated with or without BGJ398, which was injected into the mice from P7 to 1M every day. The scotopic and photopic amplitudes of a- and b-waves are shown as a function of the stimulus intensity (−4.0 to 1.0 log cd s/m$^2$ and −0.5 to 1.0 log cd s/m$^2$, respectively). Data are presented as mean ± SD. *$p < 0.05$ (unpaired t-test). n = 5 and 6 mice (vehicle-treated and BGJ398-treated, respectively).

(H, I) Effects of Ick overexpression on cilia in cells treated with a cytoplasmic dynein inhibitor. (H) NIH3T3 cells transfected with a plasmid expressing FLAG-tagged EGFP or Ick were treated with 10 μM Ciliobrevin D or DMSO for 24 h before harvest. Cells were immunostained with anti-FLAG and anti-Actub antibodies. (I) The number of cilia stained with an antibody against Actub in FLAG-positive cells was counted. Data are presented as mean ± SD. *$p < 0.05$, ****$p < 0.0001$, ns, not significant (two-way ANOVA followed by Tukey's multiple comparisons test). n = 5 experiments.

(J) Inhibition efficacy of shRNA expression constructs for *Dync2li1* knockdown. A plasmid encoding Control-shRNA, Dync2li1-shRNA2, Dync2li1-shRNA3, Dync2li1-shRNA5, or Dync2li1-shRNA6 was co-transfected with plasmids expressing a FLAG-tagged Dync2li1 and a GFP into HEK293T cells. Western blot analysis was performed using anti-FLAG and anti-GFP antibodies. GFP was used as an internal transfection control. Dync2li1-shRNA2, Dync2li1-shRNA3, Dync2li1-shRNA5, and Dync2li1-shRNA6 suppressed Dync2li1 expression.

(K, L) Effects of Ick overexpression on ciliary length in cells knocked down for *Dync2li1*. (K) A plasmid encoding Control-shRNA or Dync2li1-shRNA5 expression plasmid was co-transfected into NIH3T3 cells with or without a plasmid expressing Ick in combination with a construct encoding FLAG-tagged EGFP. Cells were immunostained with anti-FLAG and anti-Actub antibodies. (L) The length of cilia stained with an antibody against Actub in FLAG-positive cells was measured. Data are presented as mean ± SD. *$p < 0.05$, ***$p < 0.001$, ns, not significant (two-way ANOVA followed by Tukey's multiple comparisons test). Control-shRNA; Control, Control-shRNA; Ick, Dync2li1-shRNA5; Control, and Dync2li1-shRNA5; Ick, n= 79, 77, 78, and 78 cilia, respectively, from 3 experiments.

(M) Luciferase reporter gene assay using 8x Gli1-binding sites-minimal promoter-NanoLuc luciferase constructs. NIH3T3 cells were transfected with a plasmid expressing Control-shRNA, Dync2li1-shRNA3, or Dync2li1-shRNA6 along with a Nanoluc luciferase reporter construct driven by the 8x Gli1-binding sites and minimal promoter and a Firefly luciferase-expressing construct driven by the SV40 promoter and enhancer. Luciferase activities of cell lysates were measured 24 h after serum starvation with 100 nM SAG. Nanoluc luciferase activity was normalized to Firefly luciferase activity. Data are presented as mean ± SD. *$p < 0.05$, **$p < 0.01$ (one-way ANOVA followed by Tukey's multiple comparisons test). n = 3 experiments.

Nuclei were stained with DAPI. ONL, outer nuclear layer; INL, inner nuclear layer; GCL, ganglion cell layer.

**Figure S7. Generation and phenotypic analysis of *Ccrk* CKO and iCKO mice.**

(A) RT-PCR analysis of the *Ccrk* and *Bromi* transcripts in mouse tissues at 4wks. *β-actin* was used as a loading control.

(B) RT-PCR analysis of the *CCRK* and *BROMI* transcripts in the human retina. *β-actin* was used as a loading control.

(C) *In situ* hybridization analysis of *Ccrk* in the P14 mouse retina. The *Ccrk* signal was detected in the ONL.

(D) Schematic representation of the wild-type allele, targeting vector, *Ccrk* recombinant allele, *Flp* recombinant allele, and *Cre* recombinant allele. Blue and purple arrows indicate primer sets to detect the *Flp* and *Cre* recombinant alleles, respectively. Removal of exons 3 and 4 is predicted to result in a translational frameshift and loss of *Ccrk* function. Ex, exon.

(E) RT-PCR analysis of the *Ccrk* transcript in the control and *Ccrk* CKO retinas. The primers were designed within exon 2 and exon 3 of *Ccrk*. No *Ccrk* transcript was detected in the *Ccrk* CKO retina. *β-actin* was used as a loading control.

(F) Western blot analysis of lysates from HEK293T cells expressing FLAG-tagged Ccrk using anti-Ccrk and anti-FLAG antibodies. The anti-Ccrk antibody recognized the Ccrk protein.

(G) Western blot analysis of the Ccrk protein in the control and *Ccrk* CKO retinas. No Ccrk band was detected in the *Ccrk* CKO retina. α-tubulin was used as a loading control.

(H) Immunostaining of retinal sections from the control and *Ccrk* CKO mice at 1M using marker antibodies against Rhodopsin, S-opsin, and M-opsin. Severe photoreceptor degeneration was observed in the *Ccrk* CKO retina.

(I) RT-PCR analysis of the *Ccrk* transcript in the control and *Ccrk* iCKO retinas. The primers were designed within exon 2 and exon 3 of *Ccrk*. The expression level of *Ccrk* decreased in the *Ccrk* iCKO retina. *β-actin* was used as a loading control.

(J) Western blot analysis of the Ccrk protein in the control and *Ccrk* iCKO retinas. The expression level of Ccrk decreased in the *Ccrk* iCKO retina. α-tubulin was used as a loading control.

(K) Immunostaining of retinal sections from the control and *Ccrk* iCKO mice at 2M using marker antibodies against Rhodopsin, S-opsin, and M-opsin.

(L, M) ERG analysis of *Ccrk* iCKO mice. (L) Representative scotopic and photopic ERGs elicited by four different stimulus intensities (−4.0 to 1.0 log cd s/m$^2$ and −0.5 to 1.0 log cd s/m$^2$, respectively) from the control and *Ccrk* iCKO mice at 2M. (M) The scotopic and photopic amplitudes of a- and b-waves are shown as a function of the stimulus intensity. Data are presented as mean ± SD. The amplitudes of a- and b-waves were not significantly different between the control and *Ccrk* iCKO mice (unpaired t-test). n = 5 and 3 mice (control and *Ccrk* iCKO, respectively).

(N) Western blot analysis of pMak and the Mak protein in the control and *Ccrk* iCKO retinas.

Nuclei were stained with DAPI. OS, outer segment; ONL, outer nuclear layer; INL, inner nuclear layer; GCL, ganglion cell layer.

**Figure S8. Phenotypic analysis of the *Mak*$^{+/-}$; *Ccrk*$^{+/-}$ mouse retina.**

(A) Immunostaining of retinal sections from *Mak*$^{+/-}$ and *Mak*$^{+/-}$; *Ccrk*$^{+/-}$ mice at 2M using marker antibodies against Rhodopsin, S-opsin, and M-opsin. Nuclei were stained with DAPI. OS, outer segment; ONL, outer nuclear layer; INL, inner nuclear layer; GCL, ganglion cell layer.

(B, C) ERG analysis of *Mak*$^{+/-}$; *Ccrk*$^{+/-}$ mice at 2M. (B) Representative scotopic and photopic ERGs elicited by four different stimulus intensities (−4.0 to 1.0 log cd s/m$^2$ and −0.5 to 1.0 log cd s/m$^2$, respectively) from *Mak*$^{+/-}$ and *Mak*$^{+/-}$; *Ccrk*$^{+/-}$ mice. (C) The scotopic and photopic amplitudes of a- and b-waves are shown as a function of the stimulus intensity. Data are presented as mean ± SD. *$p < 0.05$ (unpaired t-test). n = 3 and

7 mice (*Mak*$^{+/-}$ and *Mak*$^{+/-}$; *Ccrk*$^{+/-}$, respectively).

(D, E) ERG analysis of *Mak*$^{-/-}$; *Ccrk*$^{+/-}$ mice at 2M. (D) Representative scotopic ERGs elicited by four different stimulus intensities (−4.0 to 1.0 log cd s/m$^2$) from *Mak*$^{-/-}$ and *Mak*$^{-/-}$; *Ccrk*$^{+/-}$ mice. (E) The scotopic amplitudes of a- and b-waves are shown as a function of the stimulus intensity. Data are presented as mean ± SD. The amplitudes of a- and b-waves were not significantly different between *Mak*$^{-/-}$ and *Mak*$^{-/-}$; *Ccrk*$^{+/-}$ mice (unpaired t-test). n = 7 and 4 mice (*Mak*$^{-/-}$ and *Mak*$^{-/-}$; *Ccrk*$^{+/-}$, respectively).

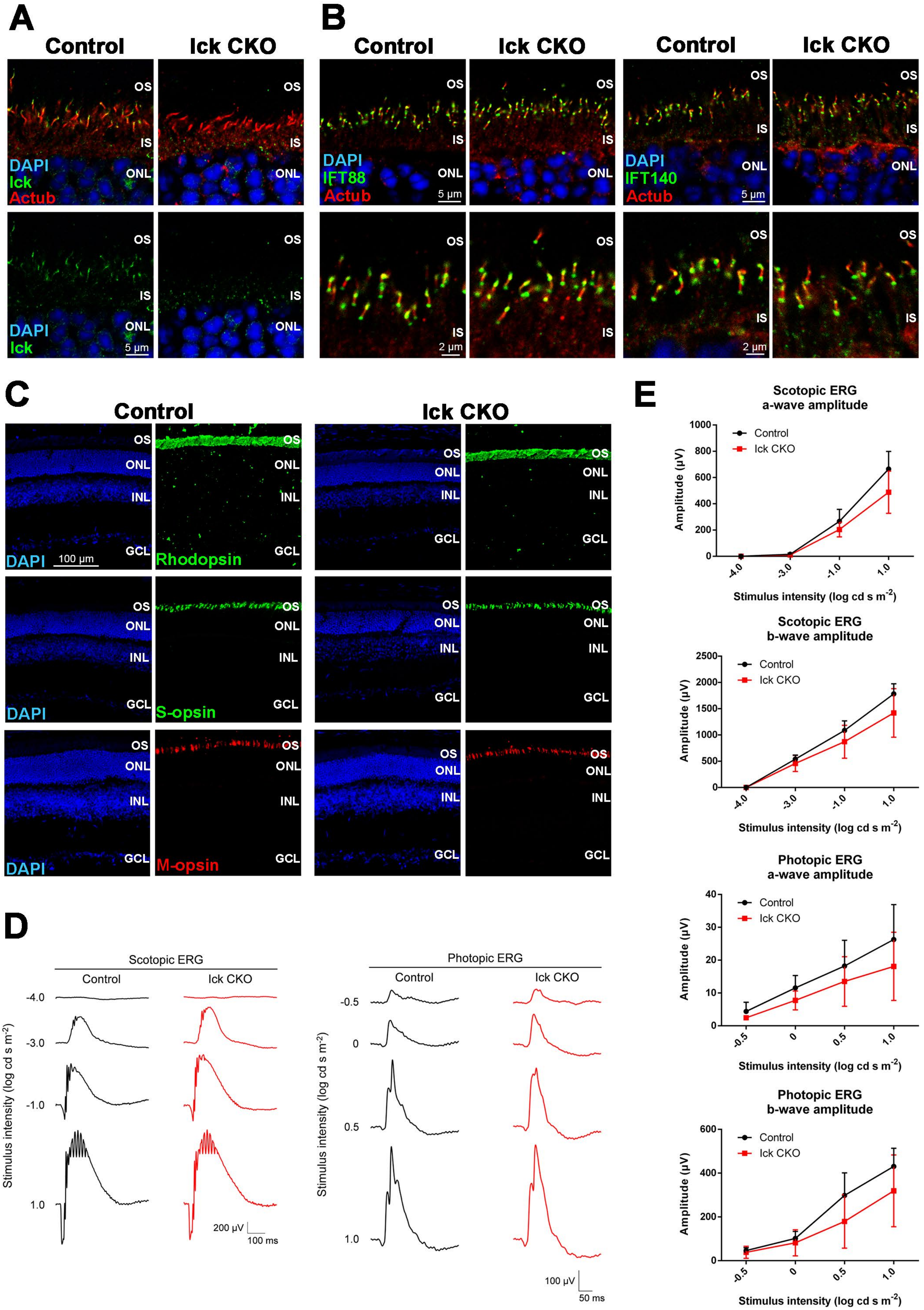
A
Control
Ick CKO
OS
IS
ONL
DAPI
Ick
Actub
5 µm
B
IFT88
IFT140
2 µm
C
INL
GCL
Rhodopsin
S-opsin
M-opsin
100 µm
D
Scotopic ERG
Photopic ERG
Stimulus intensity (log cd s m-2)
200 µV
100 ms
100 µV
50 ms
E
Scotopic ERG a-wave amplitude
Scotopic ERG b-wave amplitude
Photopic ERG a-wave amplitude
Photopic ERG b-wave amplitude
Amplitude (µV)
Stimulus intensity (log cd s m-2)

**Figure S1**

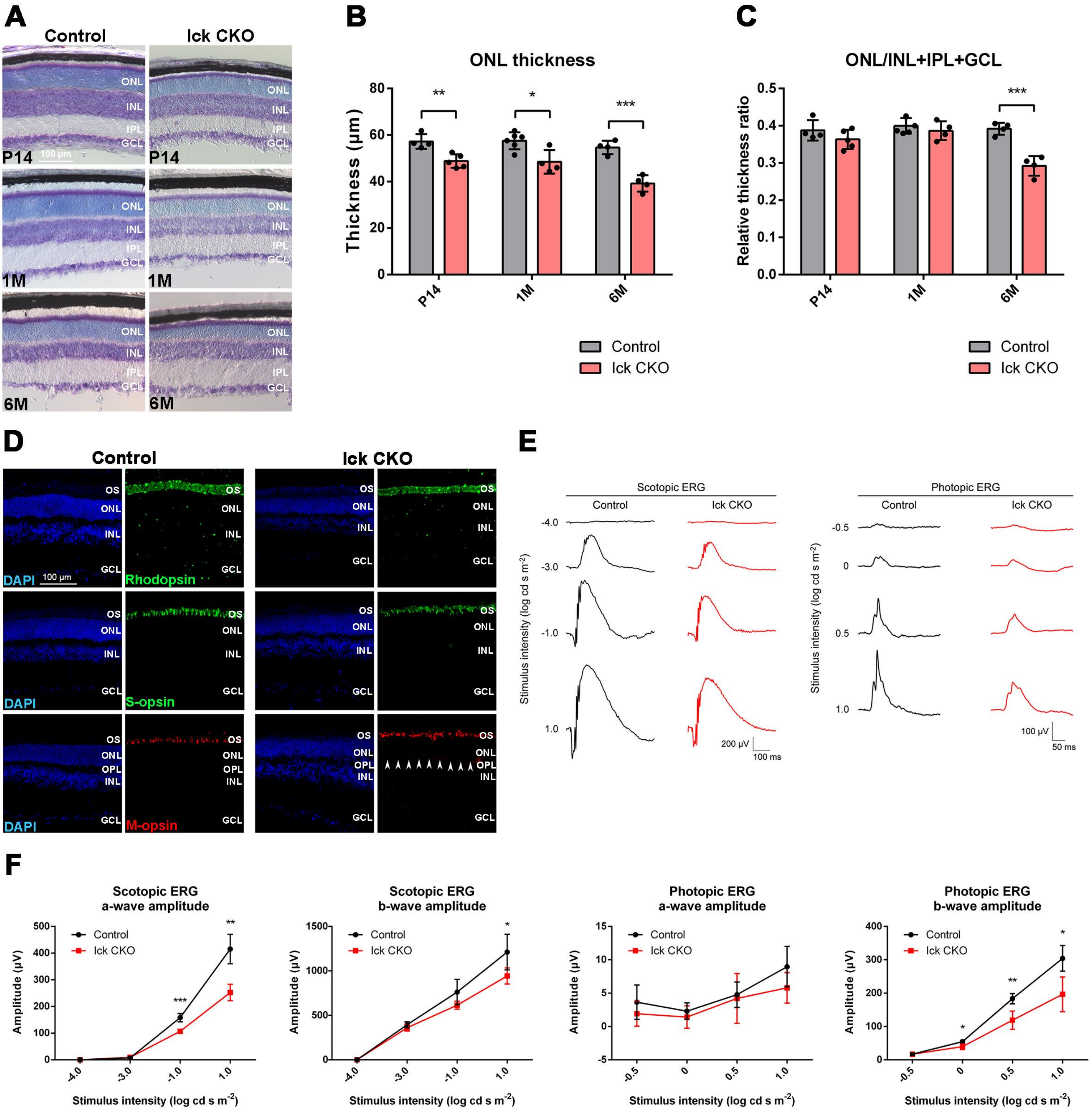

A
Control
Ick CKO
ONL
INL
IPL
GCL
P14
100 μm
1M
6M
B
ONL thickness
Thickness (μm)
P14
1M
6M
Control
Ick CKO
C
ONL/INL+IPL+GCL
Relative thickness ratio
D
Control
Ick CKO
OS
DAPI
Rhodopsin
S-opsin
M-opsin
OPL
E
Scotopic ERG
Photopic ERG
Stimulus intensity (log cd s m-2)
200 μV
100 ms
100 μV
50 ms
F
Scotopic ERG a-wave amplitude
Scotopic ERG b-wave amplitude
Photopic ERG a-wave amplitude
Photopic ERG b-wave amplitude
Amplitude (μV)
Stimulus intensity (log cd s m-2)


### Figure S2

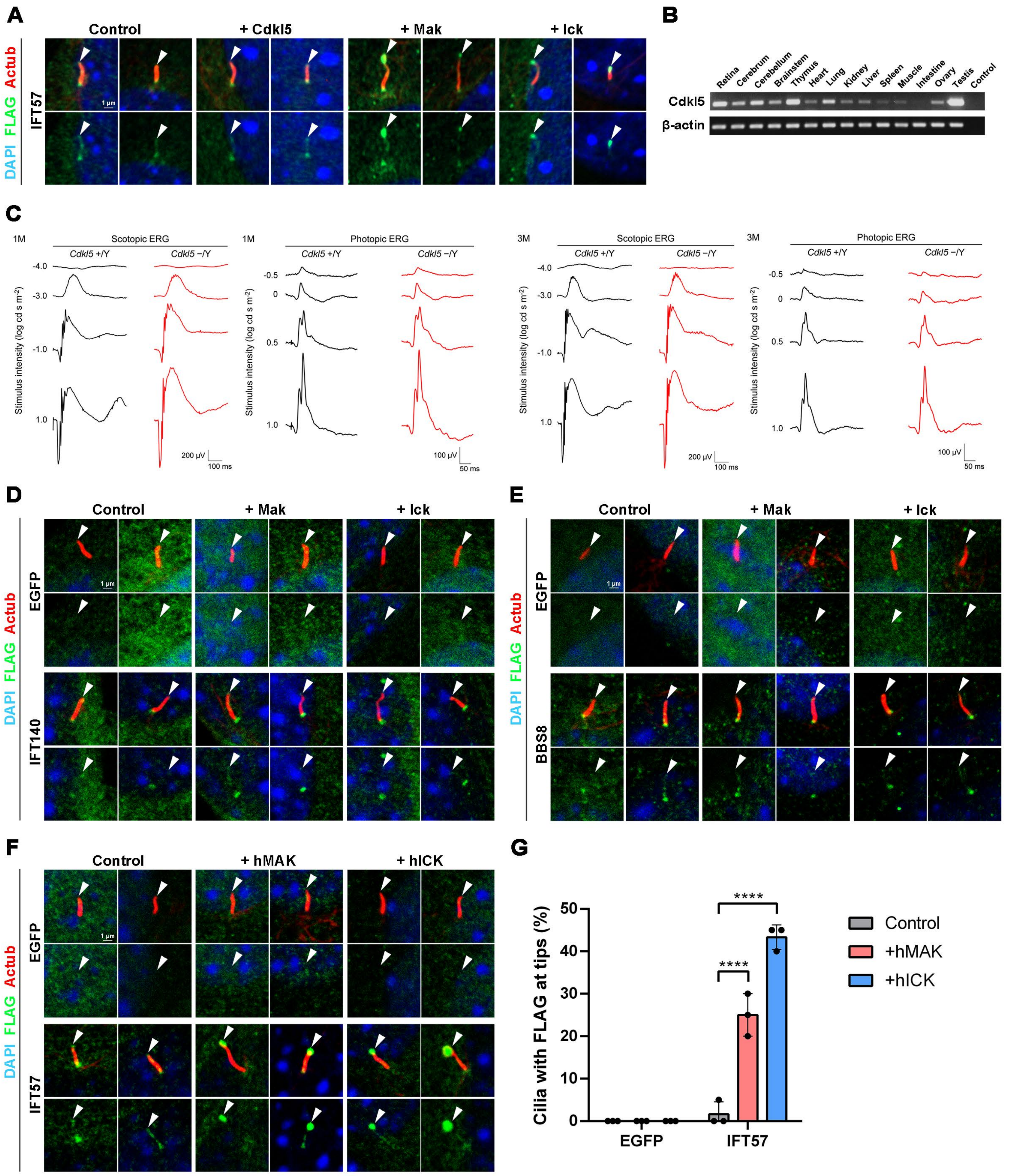
A
Control
+ Cdkl5
+ Mak
+ Ick
DAPI FLAG Actub
IFT57
1 µm
B
Retina
Cerebrum
Cerebellum
Brainstem
Thymus
Heart
Lung
Kidney
Liver
Spleen
Muscle
Intestine
Ovary
Testis
Control
Cdkl5
β-actin
C
1M
Scotopic ERG
Cdkl5 +/Y
Cdkl5 −/Y
Photopic ERG
3M
Stimulus intensity (log cd s m^-2)
-4.0
-3.0
-1.0
1.0
-0.5
0
0.5
200 µV
100 ms
100 µV
50 ms
D
Control
+ Mak
+ Ick
EGFP
IFT140
E
BBS8
F
+ hMAK
+ hICK
IFT57
G
Cilia with FLAG at tips (%)
50
40
30
20
10
0
****
EGFP
IFT57
Control
+hMAK
+hICK

### Figure S3

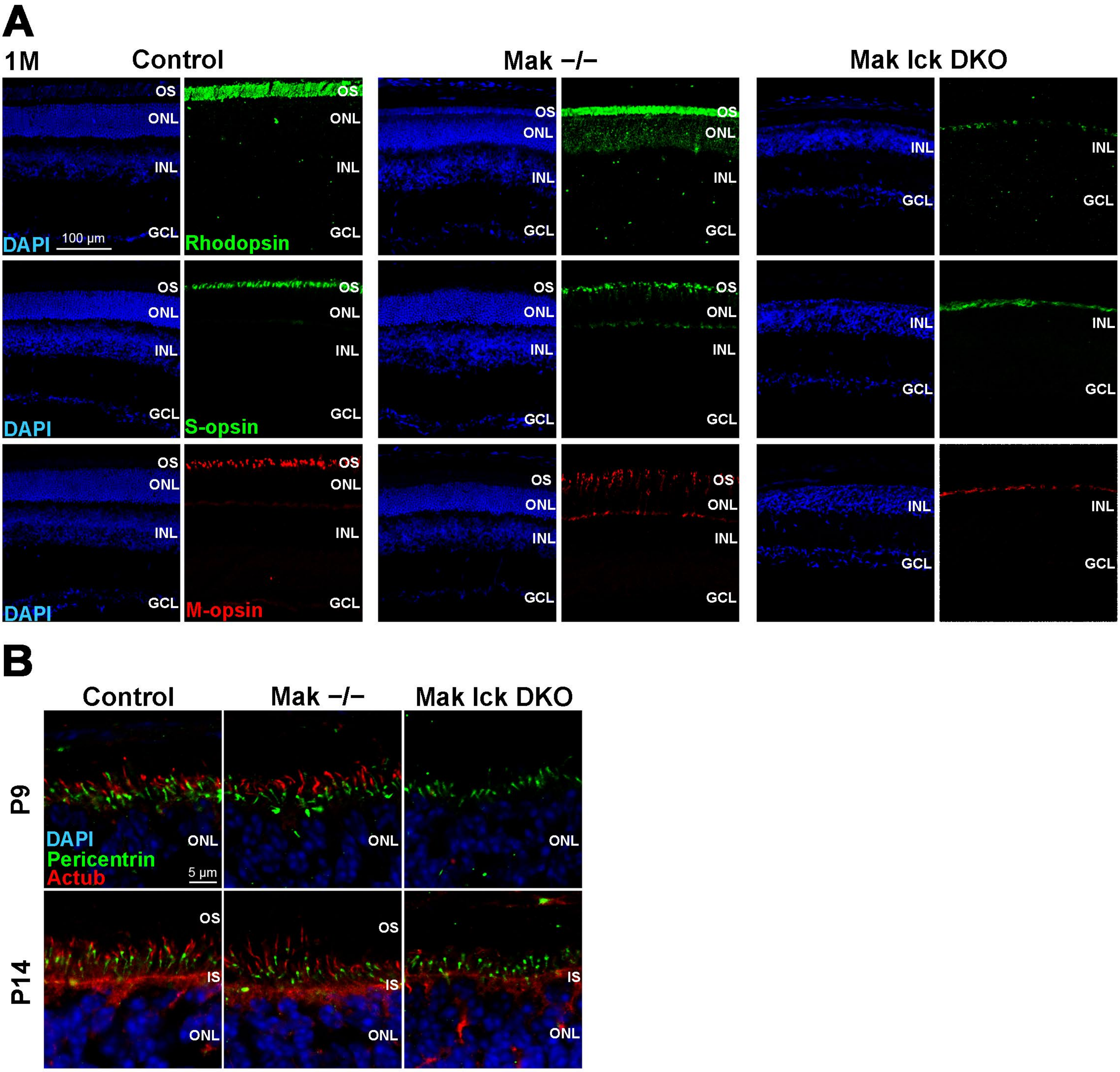
A
1M
Control
Mak −/−
Mak Ick DKO
OS
ONL
INL
GCL
DAPI
100 µm
Rhodopsin
S-opsin
M-opsin
B
P9
P14
DAPI
Pericentrin
Actub
5 µm
IS

### Figure S4

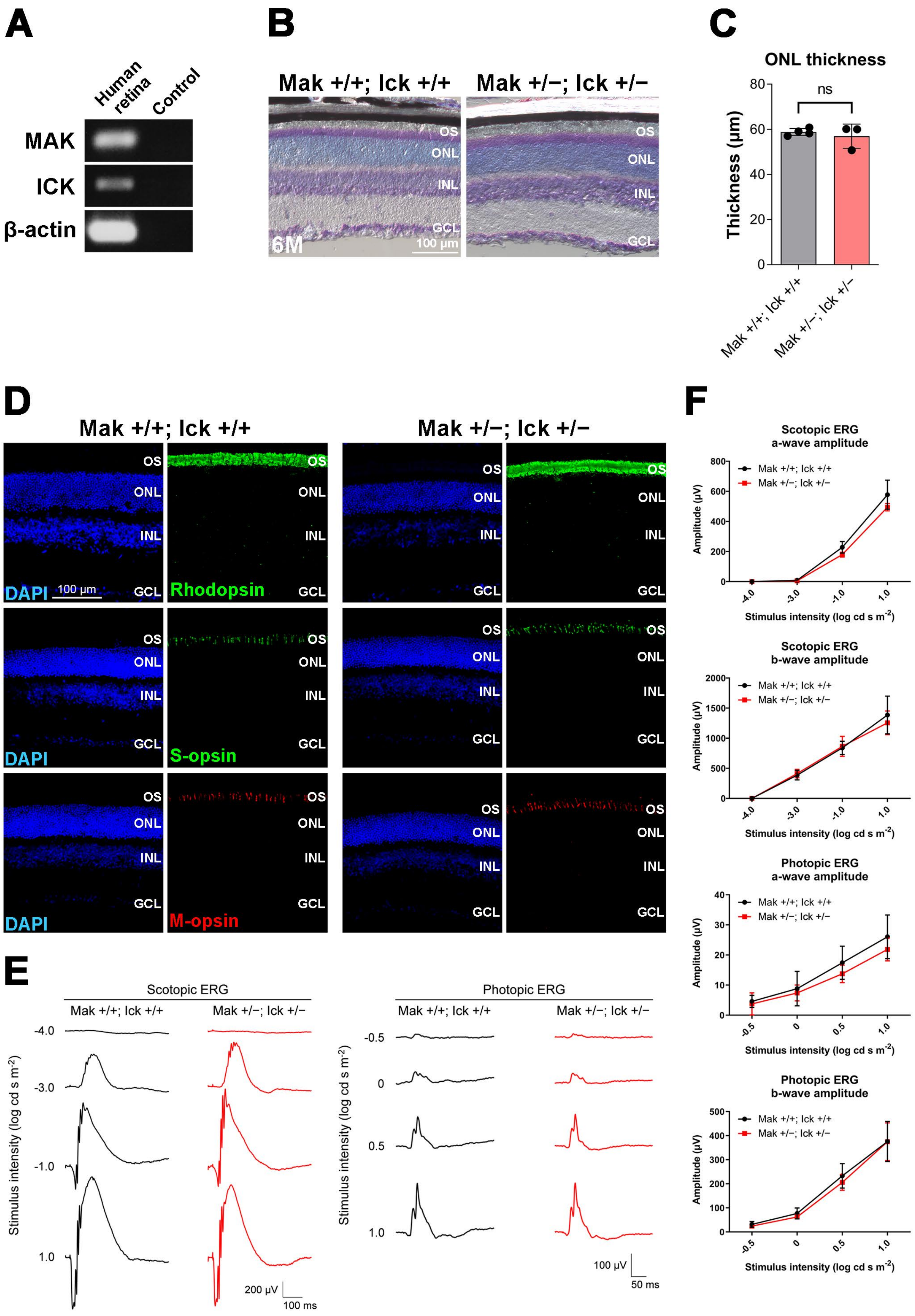
A
Human retina
Control
MAK
ICK
β-actin
B
Mak +/+; Ick +/+
Mak +/−; Ick +/−
OS
ONL
INL
GCL
6M
100 μm
C
ONL thickness
ns
Thickness (μm)
0
20
40
60
80
Mak +/+; Ick +/+
Mak +/−; Ick +/−
D
Mak +/+; Ick +/+
Mak +/−; Ick +/−
DAPI
Rhodopsin
S-opsin
M-opsin
100 μm
E
Scotopic ERG
Photopic ERG
Stimulus intensity (log cd s m-2)
-4.0
-3.0
-1.0
1.0
-0.5
0
0.5
200 μV
100 ms
100 μV
50 ms
F
Scotopic ERG a-wave amplitude
Scotopic ERG b-wave amplitude
Photopic ERG a-wave amplitude
Photopic ERG b-wave amplitude
Amplitude (μV)
Stimulus intensity (log cd s m-2)
Mak +/+; Ick +/+
Mak +/−; Ick +/−

### Figure S5

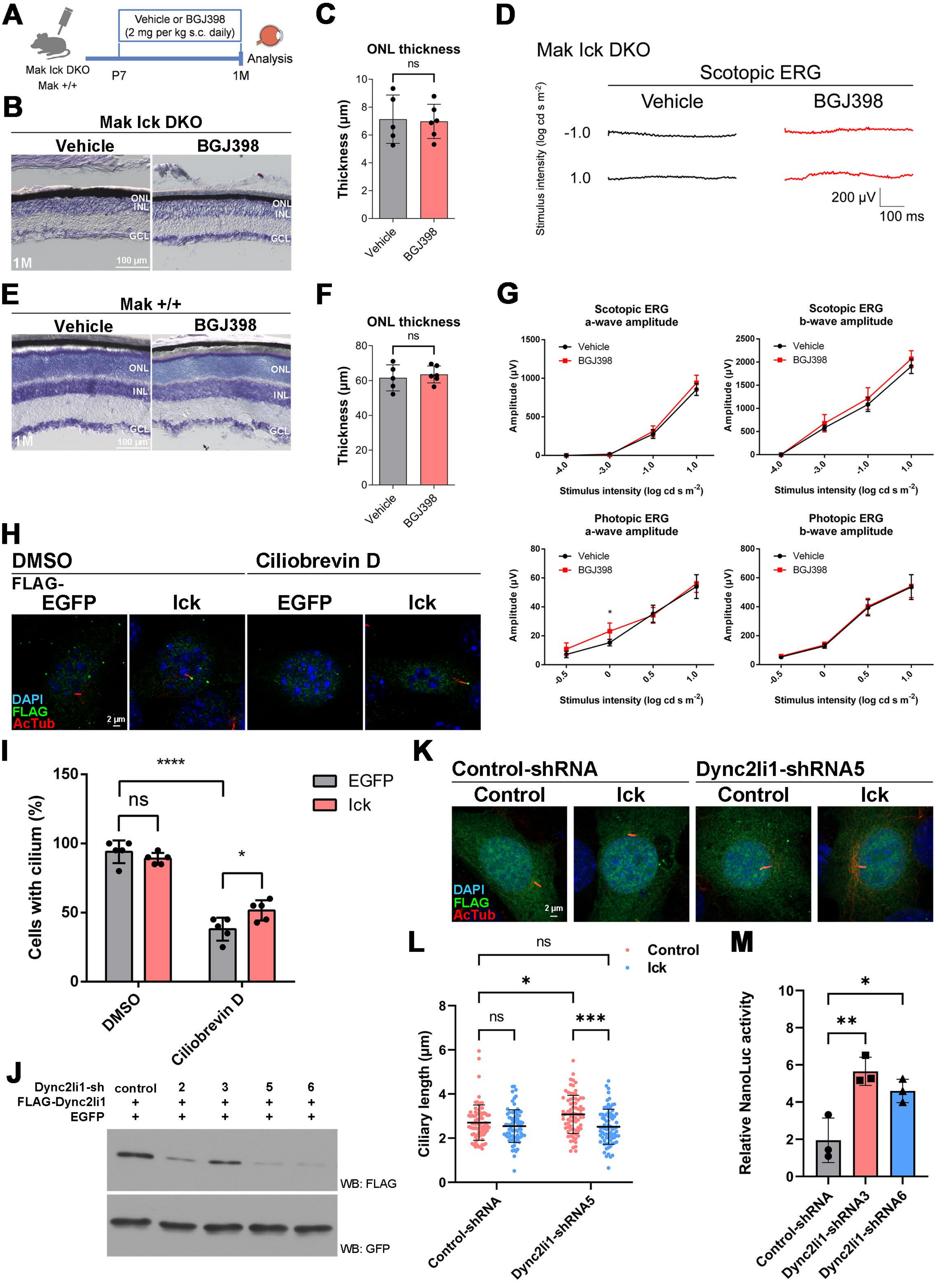

A
Vehicle or BGJ398
(2 mg per kg s.c. daily)
Mak Ick DKO
Mak +/+
P7
1M
Analysis
B
Mak Ick DKO
Vehicle
BGJ398
ONL
INL
GCL
1M
100 μm
C
ONL thickness
ns
Thickness (μm)
Vehicle
BGJ398
D
Mak Ick DKO
Scotopic ERG
Vehicle
BGJ398
Stimulus intensity (log cd s m−2)
-1.0
1.0
200 μV
100 ms
E
Mak +/+
Vehicle
BGJ398
F
ONL thickness
ns
G
Scotopic ERG a-wave amplitude
Scotopic ERG b-wave amplitude
Photopic ERG a-wave amplitude
Photopic ERG b-wave amplitude
Amplitude (μV)
Stimulus intensity (log cd s m−2)
Vehicle
BGJ398
H
DMSO
Ciliobrevin D
FLAG-
EGFP
Ick
DAPI
FLAG
AcTub
2 μm
I
Cells with cilium (%)
EGFP
Ick
ns
****
*
DMSO
Ciliobrevin D
J
Dync2li1-sh
FLAG-Dync2li1
EGFP
control
2
3
5
6
WB: FLAG
WB: GFP
K
Control-shRNA
Dync2li1-shRNA5
Control
Ick
L
Ciliary length (μm)
Control
Ick
ns
*
***
Control-shRNA
Dync2li1-shRNA5
M
Relative NanoLuc activity
**
*
Control-shRNA
Dync2li1-shRNA3
Dync2li1-shRNA6


### Figure S6

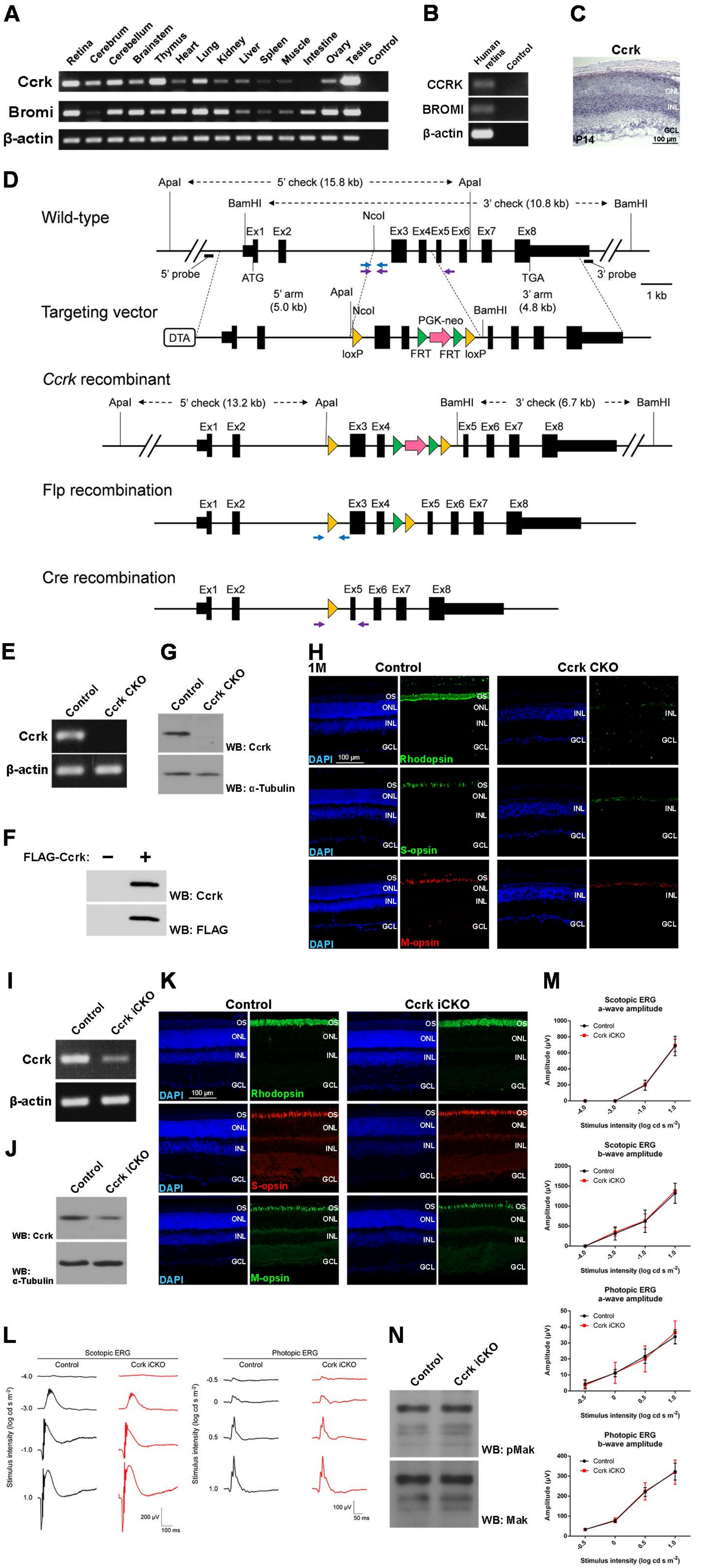
A
Retina
Cerebrum
Cerebellum
Brainstem
Thymus
Heart
Lung
Kidney
Liver
Spleen
Muscle
Intestine
Ovary
Testis
Control
Ccrk
Bromi
β-actin
B
Human retina
Control
CCRK
BROMI
β-actin
C
Ccrk
ONL
INL
GCL
P14
100 μm
D
Wild-type
Targeting vector
Ccrk recombinant
Flp recombination
Cre recombination
5' check (15.8 kb)
3' check (10.8 kb)
5' probe
3' probe
ATG
TGA
5' arm (5.0 kb)
3' arm (4.8 kb)
1 kb
DTA
PGK-neo
loxP
FRT
5' check (13.2 kb)
3' check (6.7 kb)
E
Control
Ccrk CKO
G
WB: Ccrk
WB: α-Tubulin
F
FLAG-Ccrk:
WB: FLAG
H
1M
Rhodopsin
S-opsin
M-opsin
DAPI
I
Ccrk iCKO
J
K
L
Scotopic ERG
Photopic ERG
Stimulus intensity (log cd s m⁻²)
M
Scotopic ERG a-wave amplitude
Scotopic ERG b-wave amplitude
Photopic ERG a-wave amplitude
Photopic ERG b-wave amplitude
Amplitude (μV)
N
WB: pMak
WB: Mak

### Figure S7

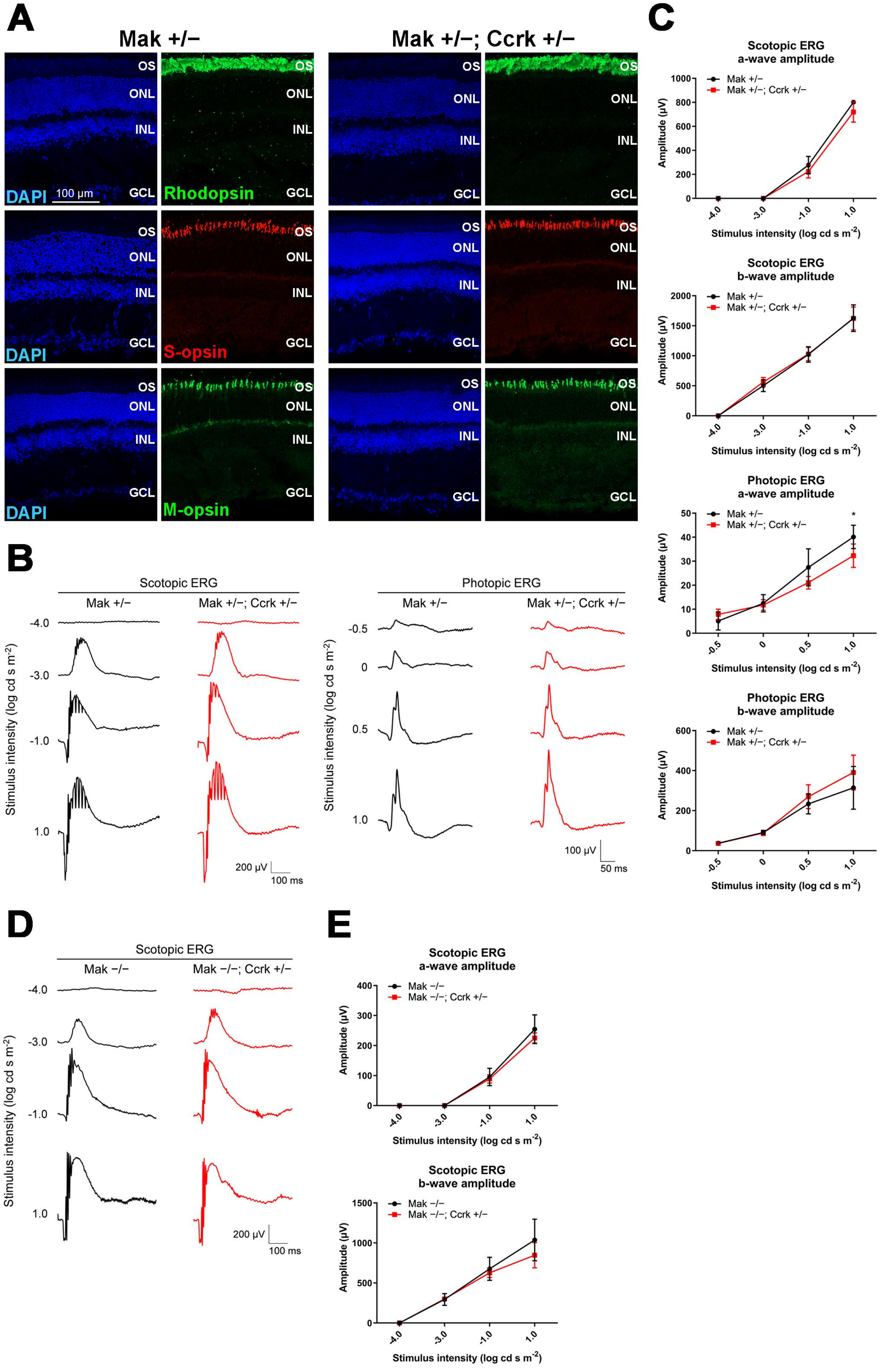
A
Mak +/−
Mak +/−; Ccrk +/−
OS
ONL
INL
GCL
DAPI
100 μm
Rhodopsin
S-opsin
M-opsin
B
Scotopic ERG
Mak +/−
Mak +/−; Ccrk +/−
-4.0
-3.0
-1.0
1.0
Stimulus intensity (log cd s m-2)
200 μV
100 ms
Photopic ERG
-0.5
0
0.5
1.0
100 μV
50 ms
C
Scotopic ERG a-wave amplitude
Scotopic ERG b-wave amplitude
Photopic ERG a-wave amplitude
Photopic ERG b-wave amplitude
Amplitude (μV)
Stimulus intensity (log cd s m-2)
*
D
Scotopic ERG
Mak −/−
Mak −/−; Ccrk +/−
E
Scotopic ERG a-wave amplitude
Scotopic ERG b-wave amplitude

### Figure S8

Table S1. Primer sequences

| Experiments | Primer name | Sequence (5' to 3') |
|---|---|---|
| Ccrk flox genotyping | CCRK-loxP-PCR-51 | CAGCAACCACATGGTGGCTCACAACCATCCA |
| | CCRK-loxP-PCR-32 | CTATTCTGTCCCGTGCTCCTGCAACCT |
| Ccrk −/− genotyping | CCRK-loxP-PCR-51 | CAGCAACCACATGGTGGCTCACAACCATCCA |
| | CCRK-loxP-PCR-32 | CTATTCTGTCCCGTGCTCCTGCAACCT |
| | CCRK-KO-PCR-31 | AGAAGCCTCTCACCTGAGACTAGCCTC |
| RT-PCR | mCdkl5-RT-PCR-51 | CACTGGTGCCACAAGAACGACATCGTC |
| | mCdkl5-RT-PCR-31 | AGATGGAAGTGGTCCTAGCACCTTCTG |
| | β-actin-F | CGTGCGTGACATCAAAGAGAA |
| | β-actin-R | TGGATGCCACAGGATTCCAT |
| | hMAK-RT-PCR-51 | GTGGGCTGTTGGAAGTATCATGGCTGA |
| | hMAK-RT-PCR-31 | TAGAGGATGCCAGCTGGTATCCTTCTG |
| | hICK-RT-PCR-51 | GGAAGAAGCATTGAGTCTGGGGAGCTG |
| | hICK-RT-PCR-31 | TTCTGGTCCCATGCAGAGGAGGTTCTC |
| | human beta-actin RT-PCR-51 | CACCACACCTTCTACAATGAGCTG |
| | human beta-actin RT-PCR-31 | GAGTCCATCACGATGCCAGTGGTA |
| | mCcrk-RT-PCR-51 | GATGGTATTCCTAACCAGGCCCTCAGA |
| | mCcrk-RT-PCR-31 | CCGATGCACAATGTTGTTGGCATGGCA |
| | mTbc1d32-RT-PCR-51 | CTGGCTATGACACGGTTGTTCAGCATG |
| | mTbc1d32-RT-PCR-31 | TGATTGAAGGAGGAGTCACTGTCGGAG |
| | hCCRK-RT-PCR-51 | CACGGCATCGTCTTCAAGGCCAAGCAC |
| | hCCRK-RT-PCR-31 | TCCACCGTGTGGGAACACAGCCTTCAG |
| | hTBC1D32-RT-PCR-51 | TCAGGGTGAAGAATGCGGCTATGATAC |
| | hTBC1D32-RT-PCR-31 | CACTATCAGAGCAATTGTCTGTGCGGT |
| In situ hybridization | mCcrk-ORF-SalI-51 | TGTCGACATGGACCAGTATTGCATCCTCGGTCGCA |
| | mCcrk-ORF-NotI-31 | TGCGGCCGCTCACCCCTCTGGGATGAAGGGCCGAATC |
| AAV titration | human_RK_promoter-QPCR-51 | TTGTCCTTCTCAGGGGAAAAGTG |
| | human_RK_promoter-QPCR-31 | TGGCAAGAAGGTGCTAGAAAAAGA |
| Construct | | |
| pCAG-FLAG-mouse Mok | mMok-ORF-XhoI-51 | TCTCGAGATGAAGAACTACAAAGCAATTGGCAAG |
| | mMok-ORF-NotI-31 | TGCGGCCGCTCAGTATTCGCCCCCTTTCCTGTTGA |
| pCAG-FLAG-mouse Cdkl1 | Cdkl1-51 | GAATTCGCCACCATGGAAAAATATGAAAAAATTGGAAAGATTGGAG |
| | Cdkl1-31 | CTCGAGAATATTTGGAAAATGGTAGTTAAATCTCTTG |
| pCAG-FLAG-mouse Cdkl2 | Cdkl2-51 | CTCGAGGCCACCATGGAGAAGTACGAGAACCTAGGATTGGT |
| | Cdkl2-31 | CTCGAGGTGTTGGTGTTCCATCCGAGGCAAATCAGC |
| pCAG-FLAG-mouse Cdkl3 | Cdkl3-51 | CTCGAGGCCACCATGGAGATGTATGAAACCCTTGGAAAAGTG |
| | Cdkl3-31 | CTCGAGAATTCTTCCCCTCACAATCGCCATCTCCA |
| pCAG-FLAG-mouse Cdkl4 | Cdkl4-51 | CTCGAGGCCACCATGGAAAAGTATGAAAAGCTAGCTAAGATC |
| | Cdkl4-31 | CTCGAGAATGTTTGGAAGATGATCGAACTTTAACTG |
| pCAG-FLAG-mouse Cdkl5 | Cdkl5-51+AAA | AAAGAATTCGCCACCATGAAGATTCCTAACATTGGTAATGTGATG |
| | Cdkl5-31+AAA | AAAGAATTCCTGTTTCCATTTCTCATGGGATGCCAAG |
| | Cdkl5-2407 | AGAGTGCCATCTCCACGACCAGACAATTCT |
| | Cdkl5-31 A+NotI+AAA | AAAGCGGCCGCACTGTTTCCATTTCTCATGGGATGCCAAG |
| pCAG-HA-mouse Cdkl5 | Cdkl5-51 EcoRI N-HA | AAAGAATTCGATGAAGATTCCTAACATTGGTAATGTGATG |
| | Cdkl5-31 NotI N-HA | AAAGCGGCCGCTCACTGTTTCCATTTCTCATGGGATGCCA |
| pCAG-FLAG-mouse Gsk3a | mGsk3a-ORF-SalI-51 | TGTCGACATGAGCGGCGGCGGGCCTTCGGGAG |
| | mGsk3a-ORF-NotI-31 | TGCGGCCGCTCAGGAAGAGCTAGCGAGGGTAGCA |
| pCAG-FLAG-mouse Gsk3b | mGsk3b-ORF-SalI-51 | TGTCGACATGTCGGGGCGACCGAGAACCACCTC |
| | mGsk3b-ORF-NotI-31 | TGCGGCCGCTCAGGTGGAGTTGGAAGCTGATGCA |
| pCAG-FLAG-mouse Dync2li1 | mDync2li1-ORF-XhoI-51 | TCTCGAGATGCCCAGTGAAACTCTCTGGGAAATC |
| | mDync2li1-ORF-NotI-31 | TGCGGCCGCTCAGGAGTCCAGCTCGATTTGCTTCCA |
| pCAG-FLAG-mouse Ccrk | mCcrk-ORF-SalI-51 | TGTCGACATGGACCAGTATTGCATCCTCGGTCGCA |
| | mCcrk-ORF-NotI-31 | TGCGGCCGCTCACCCCTCTGGGATGAAGGGCCGAATC |
| pCAG-mouse Ccrk | mCcrk-kozak-ORF-SalI-51 | GGGGTCGACGCCACCATGGACCAGTATTGCATCCTCGGTCGCA |
| | mCcrk-ORF-NotI-31 | TGCGGCCGCTCACCCCTCTGGGATGAAGGGCCGAATC |
| pCAG-HA-human MAK | hMAK-ORF-XhoI-51 | TCTCGAGATGAACCGATACACAACCATGAGACAG |
| | hMAK-ORF-NotI-31 | TGCGGCCGCCTACCGGTGGCCTCCATACTTGGCCAC |
| pCAG-FLAG-mouse Ick T157A | mIck-Y159-51-WT | GACTATGTGTCTACTAGATGGTAC |
| | mIck-T157A-31 | GGCGTACGGAGGTCTTGATCGGAT |
| pAAV-RK-FLAG-mouse Ick | mIck-ORF-ClaI(x)-infu-51 | GACAAGGACATCGAAATGAATAGATACACAACGATCAAG |
| | mIck-ORF-ClaI-infu-31 | ATCAAGCTTATCGATTCACCGCCGGGATGGGTACTTGGA |
